# Stimulation of adaptive gene amplification by origin firing under replication fork constraint

**DOI:** 10.1101/2021.03.04.433911

**Authors:** Alex J. Whale, Michelle King, Ryan M. Hull, Felix Krueger, Jonathan Houseley

## Abstract

Adaptive mutations can cause drug resistance in cancers and pathogens, and increase the tolerance of agricultural pests and diseases to chemical treatment. When and how adaptive mutations form is often hard to discern, but we have shown that adaptive copy number amplification of the copper resistance gene *CUP1* occurs in response to environmental copper due to *CUP1* transcriptional activation. Here we dissect the mechanism by which *CUP1* transcription in budding yeast stimulates copy number variation (CNV). We show that transcriptionally stimulated CNV requires TREX-2 and Mediator, such that cells lacking TREX-2 or Mediator respond normally to copper but cannot acquire increased resistance. Mediator and TREX-2 cause replication stress by tethering transcribed loci to nuclear pores, a process known as gene gating, and transcription at the *CUP1* locus causes a TREX-2-dependent accumulation of replication forks indicative of replication fork stalling. TREX-2-dependent *CUP1* gene amplification occurs by a Rad52 and Rad51-mediated homologous recombination mechanism that is enhanced by histone H3K56 acetylation and repressed by Pol32, factors known to alter the frequency of template switching during break induced replication (BIR). *CUP1* amplification is also critically dependent on late firing replication origins present in the *CUP1* repeats, and mutations that remove or inactivate these origins strongly suppress the acquisition of copper resistance. We propose that replicative stress imposed by nuclear pore association causes replication bubbles from these origins to collapse soon after firing, leaving an epigenetic scar of H3K56 acetylation that promotes template switching during later break induced replication events. The capacity for inefficient replication origins to promote copy number variation renders certain genomic regions more fragile than others, and therefore more likely to undergo adaptive evolution through *de novo* gene amplification.

## Introduction

Adaptive mutations can enable organisms to tolerate or even thrive in hostile environments. Although all kinds of mutation can be adaptive, CNV - the loss or duplication of segments of genetic material - often underlies adaptation in eukaryotic cells from fungi to mammals (1). Adaptive mutation is frequently reported in chemotherapy resistant cancers and infections (2–7), or treatment resistant animal and plant pests (8,9), so the mechanisms by which adaptive mutations form is of considerable medical, economic and societal interest.

Three major classes of mechanism are implicated in *de novo* CNV (reviewed in (10–12)): firstly non-allelic homologous recombination either in mitosis or meiosis can occur when a double strand break (DSB) forms within a region homologous to multiple sites in the genome. Strand invasion of the resected DSB into an unmatched homologue may result in duplication, deletion or translocation depending on resolution (reviewed in (13)). Secondly, non-homologous end joining can ligate unmatched DSB ends to create deletions and translocations (reviewed in (14)). Thirdly, replication fork switching between either homologous or microhomologous templates creates discontinuities in the sequence of a daughter chromatid, resulting in CNV or translocations (reviewed in (11) and (15)). All three classes can initiate further complex genome rearrangements by forming unstable species such as dicentric chromosomes or extrachromosomal DNA (reviewed in (12) and (14)), and given that adaptive mutations are normally observed only after extended selection it is often difficult to confirm formation mechanisms.

DNA replication has particular potential to invoke genetic change, and the copious CNV events induced by replication inhibitors such as hydroxyurea or aphidicolin show that template switching is a frequent outcome of replication fork stalling (16–18). Stalled forks that cannot be restarted by other means need to be repaired through recombination (19). The simplest model for recombinational repair invokes cleavage of the fork by a structure specific endonuclease, often Mus81, to create a single ended DSB that can invade the sister chromatid, a process known as Break Induced Replication (BIR) (20,21). However, this model is contested as reversal of the stalled replication fork can also form a one-ended DSB without cleavage of the template (22). Either way, replication then proceeds by a migrating D-loop rather than a standard replication fork, which has different protein components and increased risk of point mutations and template switching (11,23–28).

Adaptive mutations (including CNV) emerge through natural selection acting on random mutations. However, all types of mutation have a mechanistic cause that delimits frequency and genomic location, even if the phenotypic outcome of a given mutation is random. Mutation rate in any given genomic window may therefore be constant across time if the environment is constant or if all potentially mutagenic mechanisms acting at that locus are unaffected by environmental change. However, environmental change may disrupt normal DNA processing genome-wide or at specific genomic locations and thereby increase mutation rate. For example, induction of a gene in response to environmental change can impede oncoming replication forks, leading to site specific, environmentally stimulated mutation ((29,30) and reviewed in (31,32)).

Indeed, CNV events at the budding yeast *CUP1* locus are stimulated by transcriptional induction of the *CUP1* gene and are tightly localised to the *CUP1* region (33–35). *CUP1* encodes a metallothionein that protects yeast from environmental copper, and the copy number of the *CUP1* gene defines copper resistance such that adaptation to toxic levels of environmental copper occurs primarily through *CUP1* gene amplification (36–38). RNA polymerase II transcription can impair replication (reviewed in (39)) through head-on collisions between RNA and DNA polymerases (40), generation of torsional stresses (41,42), formation of RNA:DNA hybrids (R-loops) (29,30,43) or formation of secondary DNA structures (44,45), all of which block fork progression and/or cause replication slippage. However, *CUP1* CNV absolutely requires the histone modification Histone 3 Lysine 56 acetylation (H3K56ac) (34), which is not obviously related to any of these outcomes. Here we investigate the mechanism by which *CUP1* transcriptional induction causes CNV via H3K56ac, showing critical roles for TREX-2 and Mediator as well as late-firing replication forks adjacent to the *CUP1* genes. We propose a model involving replication origin firing, stalling and collapse that links *CUP1* transcription, replication and local histone modification to *de novo* CNV.

## Materials & Methods

### Yeast strains and media

Yeast strains used are listed in Supplementary Table 1. Deletion strains were produced by standard deletion protocols using oligonucleotides listed in Supplementary Table 2 and validated by PCR. Construction of *3xCUP1* new ARS and control strains: fragments of pRS316 containing URA3 region ± ARS were amplified using oligonucleotides listed in Supplementary Table 2 and integrated in YRH23. Construction of *3xCUP1* no ARS strain: pJH285 containing one repeat *GFP-CUP1* was formed by ligating the *Xmal Bglll-(blunt)* fragment of pFA6a-GFP-KanMX6 into pJH254 (34) digested with *XmaI EcoRV.* The 3 repeat plasmid pJH287 was formed by ligating 3 fragments - pJH285 *ClaI SalI,* pJH285 *XhoI BglII* and pJH285 *Bam*HI *Eco*RI – simultaneously into *EcoRI ClaI* digested pJH264 (34). This construct was integrated into genome of YRH15 as described in (34).

All cells were cultured in shaking incubators at 30°C, 200 rpm. Overnight cultures and cells used in standard experiments were grown in yeast nitrogen base (YNB) media (which contains 250 nM CuSO_4_) that was supplemented with CSM amino acids and 2% glucose (or 2% raffinose with 0.02% galactose when stated). YNB media and supplements were purchased from Formedium. Pre-cultures for all copper experiments were grown to saturation (~2 days) in 4 ml cultures of YNB media and then diluted 1:2000 for subsequent treatments. For Nicotinamide treatment, cells were cultured for one week in 4 ml YNB media with a final concentration of 5 mM Nicotinamide (Sigma I17451). For copper treatment, cells were grown in 4 ml YNB media ± 0.3 mM CuSO_4_ for one week. For Northern blot analysis, cells were grown in 4 ml YNB media with 2% glucose for 6 hours, diluted and grown overnight in 25 ml same media to 0.6-0.8×10^7^ cells/ml. Un-induced cells were harvested, cells were diluted to 0.15×10^7^ cells/ml in 25 ml with 0.3 mM CuSO_4_ and grown for 6 hours before harvesting 2×10^7^ cells by centrifugation and freezing on N2. For TrAEL-seq experiments, cells were pre-cultured by inoculation in 4 ml yeast peptone broth containing 2% Raffinose (YP Raf) for ~6 h at 30°C with shaking at 200 rpm. These cells were then diluted in 100 ml YP Raf (wild-type and *rad52Δ* cells ~1:500, *sac3Δ* cells ~1:50) and growth continued at 30°C 200 rpm for ~16 hours until OD_600_ reached ~0.2. These 100 ml cultures were then split, with 50 ml transferred into 50 ml YP Raf media and the other 50 ml being transferred into 50 ml YP Raf media containing 0.02% galactose for 6 hours at 30°C 200 rpm. Cells were centrifuged 1 min at 4,600 rpm, resuspended in 70% ethanol at 1×10^7^ cells/ml and stored at −70°C. YP media, raffinose and galactose were purchased from Formedium.

### Adaptation Assay

From saturated cultures grown ± 0.3 mM CuSO_4_, a 1:80 dilution in 200 μl of YNB media was placed in every well in a flat-bottomed 96-well cell culture plate with CuSO_4_ at the required concentration. Plates were sealed using a gas-permeable membrane and incubated at 30°C with shaking for 3 days. Cells were resuspended and OD660 was measured by a BD FLUOstar Omega plate reader. Area-Under-Curve for plots of OD660 against [CuSO_4_] were calculated for each sample and compared by one way ANOVA with Sidak’s multiple comparison correction in GraphPad Prism (v8.2.1).

### DNA extraction and Southern blotting

From a saturated culture, 2 ml of cells were washed in 50 mM EDTA and then spheroplasted using 250 μl of 0.34 U/ml lyticase (Sigma L4025) in 1.2 M sorbitol, 50 mM EDTA and 10 mM DTT at 37°C for 45 minutes. These cells were centrifuged at 1000 rcf, gently resuspended in 400 μl of 100 μg/ml RNase A (Sigma R4875), 50 mM EDTA and 0.3% SDS, and then incubated for 30 minutes at 37°C. After this, 4 ul of 20 mg/ml proteinase K (Roche 3115801) was added, mixed by inverting the samples and heated at 65°C for 30 minutes. The samples were then left to cool to room temperature before adding 160 μl of 5 M KOAc, then being mixed by inversion and chilled on ice for 1 hour. These samples were centrifuged at 20,000 rcf for 10 minutes before the supernatant was poured into a new tube containing 500 μl of phenol:chloroform (pH8) and placed on a rotating wheel for 30 minutes. After centrifugation at 10,000 rcf for 10 minutes, the upper phase was extracted using wide bore pipette tips and precipitated in 400 μl isopropanol. Pellets were then washed in 70% ethanol, left to air-dry and then digested overnight at 37°C in 50 μl TE with 20 U *Eco*RI-HF (NEB). Samples were extracted with 50 μl phenol:chloroform, then ethanol precipitated in a 1.5 ml tube containing 112.5 μl 100% ethanol and 4.5 μl 3 M NaOAc before centrifugation at 20,000 rcf for 15 minutes. After washing in 70% ethanol, the pellets were dissolved for 1 hour in 20 μl TE. Loading dye was added and samples were separated on 25 cm 0.8% or 1% TBE gels at 120 V for 16.5 hours. Gels were denatured in 0.25 N HCl for 15 minutes, neutralised in 0.5 N NaOH for 45 minutes and washed twice in 1.5 M NaCl, 0.5 M Tris (pH7.5) for 20 minutes each. Samples were transferred to HyBond N+ membrane in 6x SSC through capillary action overnight and fixed by UV crosslinking using a Stratagene UV Stratlinker. Membranes were probed using random primed probes (listed in Supplementary Table 2) in 10ml UltraHyb (AM8669 ThermoFisher Scientific) at 42°C then washed with 0.1x SSC 0.1% SDS. Quantification of Southern bands was performed using ImageQuant (Version 7.0, GE), and CNV calculated as (intensity of all CNV bands / intensity of CNV and parental bands) × 100. Statistical analysis of CNV levels was performed by one-way ANOVA with Sidak’s multiple comparison correction in GraphPad Prism (v8.2.1).

### RNA extraction and northern blotting

Frozen cell pellets were lysed by 5 minutes vortexing at 4°C with ~50 μl glass beads and 40 μl GTC-phenol (2.1 M guanidine thiocyanate, 26.5 mM Na citrate pH7, 5.3 mM EDTA, 76 mM β-mercaptoethanol, 1.06% N-lauryl sarcosine, 50% phenol pH7). 600 μl GTC-phenol was added, mixed, and samples were heated at 65°C for 10 minutes then placed on ice for 10 minutes. 160 μl 100mM NaOAc pH 5.2 and 300 μl chloroform:isoamyl alcohol (24:1) were added, samples were vortexed and centrifuged at top speed for 5min. The upper phase was re-extracted first with 500 μl phenol:chloroform pH 7 (1:1) and then with 500 μl chloroform:isoamyl alcohol (24:1) before precipitation with 1 ml ethanol. Pellet was washed with 70% ethanol, re-suspended in 6 μl water and quantified using Quant-IT RiboGreen (ThermoFisher, R11490). 1 μg RNA was resolved per lane on 1.2% glyoxal agarose gels, blotted and probed with a random primed probe against *CUP1* ORF (Supplementary Table 4) as described (46).

### Candidate Genetic Screen

A total of 206 strains from the Yeast Deletion Collection (Invitrogen 95401.H2) and other sources were streaked out on YPD agar plates and then restreaked for single colonies on YPD plates containing 300 μg/ml G418. Pre-cultures were grown to saturation in 4 ml YNB media at 30°C, diluted 1:2000 in 4 ml YNB media with and without 5 mM Nicotinamide and then incubated for one week at 30°C with shaking. DNA extraction and Southern blotting was performed as described above. CNV rates of mutants were obtained by comparing the percentage of CNV alleles in nicotinamide-treated mutants to nicotinamide-treated wildtype cells, and calculating the fold change in CNV.

For Network analysis, factors from the CNV screen and their first neighbours were imported into Cytoscape (v3.7.2) using stringApp (v1.6.0) (47) to retrieve *S. cerevisiae* protein interaction data and to construct the network. The Edge-weighted Spring Embedded layout was applied using “stringdb score” to determine edge length, and node size and colour were mapped to fold-change in CNV from the CNV screen with labels applied to the strongest CNV enhancers and suppressors (>2 or <0.5 fold-change in CNV respectively). Cluster Analysis was performed on this network using the ClusterONE app (v1.0) (48) which was used to identify clusters of proteins, and clusters with a similar impact on fold-change in CNV have their functional categories displayed.

### TrAEL-seq library preparation and sequencing

1-3×10^7 cells fixed in ethanol were pelleted by centrifugation for 30 s at 20,000 g, rinsed in 1 ml PFGE wash buffer (10 mM Tris HCl pH 7.5, 50 mM EDTA), and resuspended in 60 μl PFGE wash buffer containing 1 μl lyticase (17 U/μl 10 mM KPO4 pH 7, 50% glycerol Merck L2524 >2000 U/mg) then incubated for 10 min at 50°C. 40 μl of molten CleanCut agarose (Bio-Rad 1703594) cooled to 50°C was added, samples were vortexed vigorously for 5 s and pipetted into a plug mould (Bio-RAD 1703713), then left to solidify for 30 min at 40°C. Plugs were transferred into a 2 ml Eppendorf that contained 500 μl PFGE wash buffer containing 10 μl 17 U/ml lyticase and left to incubate at 37°C for 1 hour. This solution was removed and replaced with 500 μl PK buffer (100 mM EDTA pH 8, 1 mg/ml Proteinase K, 1% sodium N-lauroyl sarcosine, 0.2% sodium deoxycholate) at 50°C overnight. The plugs were then rinsed in 1 ml TE and washed with 1 ml TE for 1 hour with rocking. Plugs were then washed twice with 1 ml TE containing 10 mM PMSF (Merck 93482) for 1 hour with rocking. Finally, plugs were digested in 200 μl TE containing 1 μl 1000U/ml RNase T1 (Thermo EN0541) at 37°C for 1 hour before being stored at 4°C in 1ml TE.

A ½ plug was used for each sample (referred to here on in as plugs). Plugs were equilibrated once in 100 μl 1x TdT buffer (NEB) for 30 min at room temperature, then incubated for 2 h at 37°C in 100 μl 1x TdT buffer containing 4 μl 10 mM ATP and 1 μl Terminal Transferase (NEB M0315L). Plugs were rinsed with 1 ml tris buffer (10 mM Tris HCl pH 8.0), equilibrated in 100 μl 1x T4 RNA ligase buffer (NEB) containing 40 μl 50% PEG 8000 for 1 hour at room temperature then incubated overnight at 25°C in 100 μl 1x T4 RNA ligase buffer (NEB) containing 40 μl 50% PEG 8000, 1 μl 10 pM/μl TrAEL-seq adaptor 1 (49) and 1 μl T4 RNA ligase 2 truncated KQ (NEB M0373L). Plugs were then rinsed with 1 ml tris buffer, transferred to 15 ml tubes and washed three times in 10 ml tris buffer with rocking at room temperature for 1-2 hours each, then washed again overnight under the same conditions. Plugs were equilibrated for 15 min with 1 ml agarase buffer (10 mM Bis-Tris-HCl, 1 mM EDTA pH 6.5), then the supernatant removed and 50 μl agarase buffer added. Plugs were melted for 20 min at 65°C, transferred for 5 min to a heating block pre-heated to 42°C, 1 μl β-agarase (NEB M0392S) was added and mixed by flicking without allowing sample to cool, and incubation continued at 42°C for 1 h. DNA was ethanol precipitated with 25 μl 10 M NH4OAc, 1 μl GlycoBlue, 330 μl of ethanol and resuspended in 10 μl 0.1x TE. 40 μl reaction mix containing 5 μl Isothermal amplification buffer (NEB), 3 μl 100 mM MgSO4, 2 μl 10 mM dNTPs and 1 μl Bst 2 WarmStart DNA polymerase (NEB M0538S) was added and sample incubated 30 min at 65°C before precipitation with 12.5 μl 10 M NH4OAc, 1 μl GlycoBlue, 160 μl ethanol and re-dissolving pellet in 130 μl 1x TE. The DNA was transferred to an AFA microTUBE (Covaris 520045) and fragmented in a Covaris E220 using duty factor 10, PIP 175, Cycles 200, Temp 11°C, then transferred to a 1.5 ml tube containing 8 μl pre-washed Dynabeads MyOne streptavidin C1 beads (Thermo, 65001) re-suspended in 300 μl 2x TN (10 mM Tris pH 8, 2 M NaCl) along with 170 μl water (total volume 600 μl) and incubated 30 min at room temperature on a rotating wheel. Beads were washed once with 500 μl 5 mM Tris pH 8, 0.5 mM EDTA, 1 M NaCl, 5 min on wheel and once with 500 μl 0.1x TE, 5 min on wheel before re-suspension in 25 μl 0.1x TE. Second end processing and library amplification were performed with components of the NEBNext Ultra II DNA kit (NEB E7645S) and a NEBNext Multiplex Oligos set (e.g. NEB E7335S). 3.5 μl NEBNext Ultra II End Prep buffer, 1 μl 1 ng/μl sonicated salmon sperm DNA (this is used as a carrier) and 1.5 μl NEBNext Ultra II End Prep enzyme were added and reaction incubated 30 min at room temperature and 30 min at 65°C. After cooling, 1.25 μl 10 pM/μl TrAEL-seq adaptor 2 (49), 0.5 μl NEBNext ligation enhancer and 15 μl NEBNext Ultra II ligation mix were added and incubated 30 min at room temperature. The reaction mix was removed and discarded and beads were rinsed with 500 μl wash buffer (5 mM Tris pH 8, 0.5 mM EDTA, 1 M NaCl) then washed twice with 1 ml wash buffer for 10 min on wheel at room temperature and once for 10 min with 1 ml 0.1x TE. Libraries were eluted from beads with 11 μl 1x TE and 1.5 μl USER enzyme (NEB) for 15 min at 37°C, then again with 10.5 μl 1x TE and 1.5 μl USER enzyme (NEB) for 15 min at 37°C, and the two eluates combined. An initial test amplification was used to determine the optimal cycle number for each library. For this, 1.25 μl library was amplified in 10 μl total volume with 0.4 μl each of the NEBNext Universal and any NEBNext Index primers with 5 μl NEBNext Ultra II Q5 PCR master mix. Cycling program: 98°C 30s then 18 cycles of (98°C 10 s, 65°C 75 s), 65°C 5 min. Test PCR was cleaned with 8 μl AMPure XP beads (Beckman A63881) and eluted with 2.5 μl 0.1x TE, of which 1 μl was examined on a Bioanalyser high sensitivity DNA chip (Agilent 5067-4626). Ideal cycle number should bring final library to final concentration of 1-3 nM, noting that the final library will be 2-3 cycles more concentrated than the test anyway. 21 μl of library was then amplified with 2 μl each of NEBNext Universal and chosen Index primer and 25 μl NEBNext Ultra II Q5 PCR master mix using same conditions as above for calculated cycle number. Amplified library was cleaned with 40 μl AMPure XP beads (Beckman A63881) and eluted with 26 μl 0.1x TE, then 25 μl of this was again purified with 20 μl AMPure XP beads and eluted with 11 μl 0.1x TE. Final libraries were quality controlled and quantified by Bioanalyser (Agilent 5067-4626) and KAPA qPCR (Roche KK4835). Libraries were sequenced on an Illumina NextSeq 500 as High Output 75 bp Single End by the Babraham Institute Next Generation Sequencing facility.

### TrAEL-seq data processing

Unique Molecular Identifier (UMI) deduplication and mapping: Scripts used for UMI-handling as well as more detailed information on the processing are available here: https://github.com/FelixKrueger/TrAEL-seq). Briefly, TrAEL-seq reads are supposed to carry an 8 bp in-line barcode (UMI) at the 5’-end, followed by a variable number of 1-3 thymines (T). Read structure is therefore NNNNNNNN(T)nSEQUENCESPECIFIC, where NNNNNNNN is the UMI, and (T)n is the poly(T). The script TrAELseq_preprocessing.py removes the first 8bp (UMI) of a read and adds the UMI sequence to the end of the readID. After this, up to 3 T (inclusive) at the start of the sequence are removed. Following this UMI and Poly-T pre-processing, reads underwent adapter- and quality trimming using Trim Galore (v0.6.5; default parameters; https://github.com/FelixKrueger/TrimGalore). UMI-pre-processed and adapter-/quality trimmed files were then aligned to the respective genome using Bowtie2 (v2.4.1; option: ╌local; http://bowtie-bio.sourceforge.net/bowtie2/index.shtml) using local alignments. Finally, alignment results files were deduplicated using UmiBam (v0.2.0; https://github.com/FelixKrueger/Umi-Grinder). This procedure deduplicates alignments based on the mapping position, read orientation as well as the UMI sequence.

To assist with the interpretation of aligned multi-copy sequences, the *P_GAL1_-HA cup1* samples were treated in a more specialised way before entering the TrAEL-seq processing procedure outlined above: Prior to TrAEL-seq pre-processing, sequences were deduplicated based on the first 23bp on their 5’-end (using the script TrAELseq_sequence_based_deduplication.py). This region contains both the UMI sequence as well as the first 15bp of genomic sequence, and should thus help identify (and remove) PCR amplified multi-copy sequences that would under normal conditions survive the UMI-aware deduplication procedure by aligning to several different genomic regions at random. Following deduplication-by-sequence and TrAEL-seq pre-processing, these sequences were aligned to a modified version of the yeast genome containing the additional Pgal-HA control sequences. To avoid multi-mapping artefacts arising from the integration of these sequences, the following two stretches of genomic sequence were masked by Ns: a) *CUP1* (chromosome VIII:212266-216251), and b) *P_GAL1_*. (chromosome II:278352-279023). Reads were then trimmed and mapped as above.

De-duplicated mapped reads were imported into SeqMonk v1.47 (https://www.bioinformatics.babraham.ac.uk/projects/seqmonk/) and immediately truncated to 1 nucleotide at the 5’ end, representing the last nucleotide 5’ of the strand break. Reads were then summed in running windows as described in figure legends. Windows overlapping with non-single copy regions of the genome were filtered (rDNA, 2μ, mtDNA, sub-telomeric regions, Ty elements and LTRs), and total read counts across all included windows were normalised to reads per million mapped. A further Enrichment Normalisation (20-90%) was applied to match the read count distributions of the *P_GAL1_-HA cup1* libraries. Read counts were exported for the consensus *P_GAL1_-HA cup1* region and plotted in GraphPad Prism 8. Comparison of datasets was performed using edgeR implemented in SeqMonk (50). For read polarity plots, forward and reverse read counts were quantitated in running windows as specified in the relevant figure legends before export for plotting using R v4.0.0 in RStudio. Read polarity values were calculated and plotted as either dots (individual samples) or as a continuous line (multiple sample display) for each quantification window using the formula read polarity = (R-F)/(R + F), where F and R relate to the total forward and reverse read counts respectively. The R code to generate these plots can also be found here: https://github.com/FelixKrueger/TrAEL-seq.

### Data Availability and Image processing

Gel images were processed in ImageQuant TL (v7.0) which involved cropping, rotating and altering contrast of whole images to improve visualisation of bands.

Sequencing data is available on GEO accession number GSE165163

## Results

### *A genetic screen identifies enhancers and suppressors of* CUP1 *CNV*

The *CUP1* locus on chromosome VIII is composed of 1 or more tandem copies of a 2 kb sequence containing the *CUP1* gene with a copper sensitive *CUP1* promoter and a poorly defined replication origin (an autonomously replicating sequence or ARS) (Figure 1A). To identify factors important for *CUP1* CNV, we performed a candidate genetic screen using mutants from the Yeast Deletion Collection (51). The genetic background of this collection includes a 13 copy *CUP1* array that rarely amplifies but does undergo transcriptionally stimulated contraction through an H3K56ac-dependent mechanism (34). Nicotinamide, an inhibitor of H3K56 deacetylases, accentuates CNV stimulation by transcription, such that even basal *CUP1* expression in the absence of copper becomes sufficient to cause measurable contractions (34). Fold-change in CNV relative to wild type can then be quantified by Southern blot after 10 generations ± nicotinamide to reveal enhancers or suppressors of CNV such as *MRC1* or *CTF4* (Figure 1B)(see Supplementary Table 3 for data on all strains tested).

**Figure 1.**
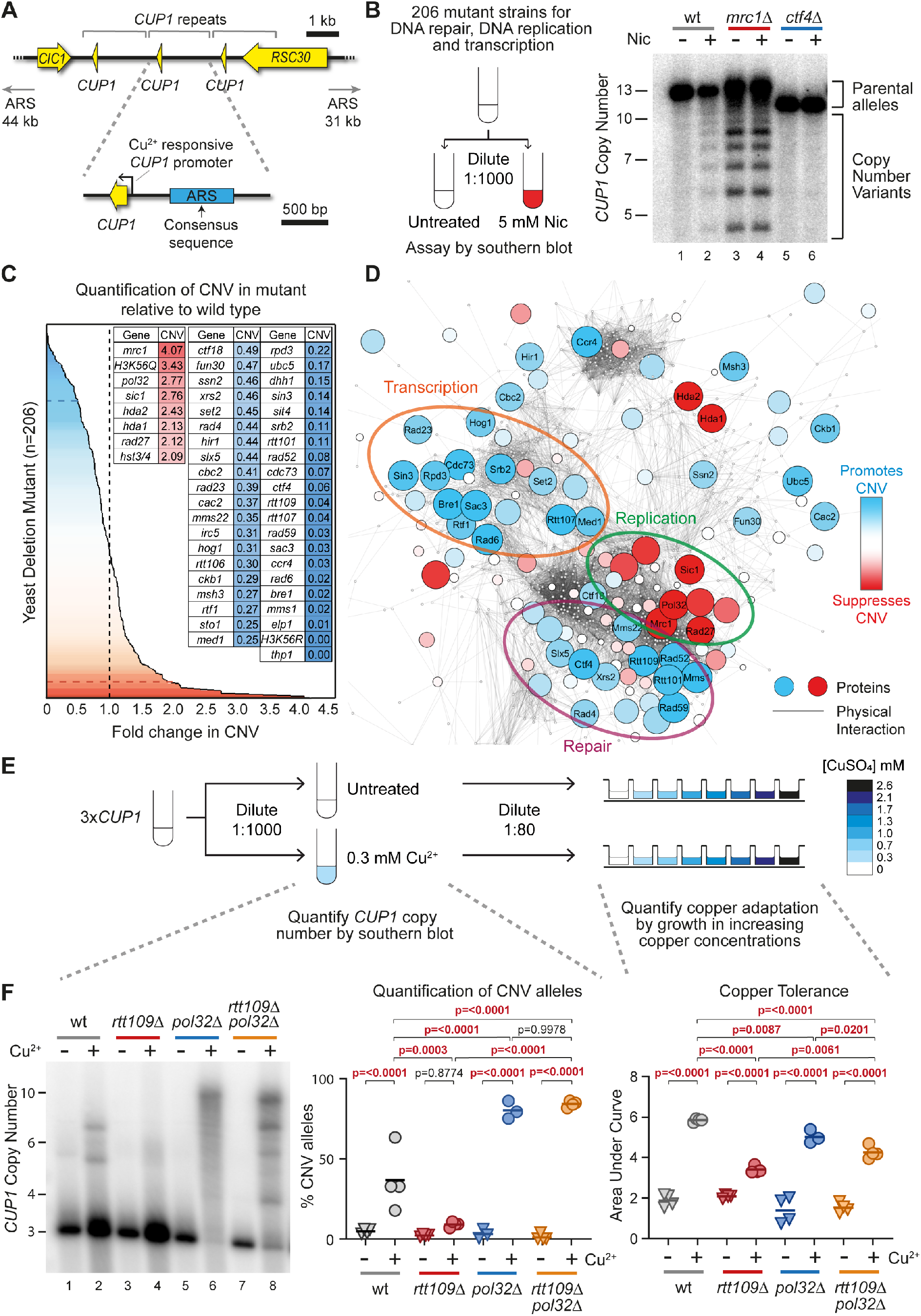
A screen for genes regulating transcriptionally stimulated CNV. **(A)** Schematic of the *CUP1* array and the surrounding region of Chromosome VIII. ORFs are shown in yellow, and grey brackets indicate the repeated region. Each repeat contains a replication origin (ARS); the ARS consensus sequence was defined in (116) and the blue rectangle denotes a sequence with ARS activity experimentally validated in (98) though the actual limits of the ARS element have not been determined. Schematic shows cells with 3 copies of the *CUP1* gene although the BY4741 background used as in the initial screen contains 13 *CUP1* copies. The nearest adjacent replication origins (ARS elements) in both directions are indicated. **(B)** Schematic of the candidate genetic screen for regulators of CNV and a representative Southern blot used for quantification of CNV. Wild type (wt) and indicated mutant cells from yeast haploid deletion collection (Invitrogen 95401.H2) and other sources (Supplementary Table 3) were grown to saturation then diluted 1:1000 and re-grown to saturation (10 generations) ± 5 mM Nicotinamide (Nic) and subject to southern analysis of *CUP1* copy number. Quantification of % CNV alleles was calculated as the percentage of alleles deviating from the parental copy number and fold-change was calculated relative to % CNV alleles in nicotinamide-treated wild-type cells. **(C)** Summary of genetic screen plotting fold-change in nicotinamide-induced CNV of 206 deletion strains relative to wild-type. Grey dashed line indicates wild-type fold-change in CNV. The blue dashed line represents cut-off for CNV-suppressing mutations with a <0.5 fold change in CNV and red dashed lines indicates the cut-off for CNV enhancing mutations with a >2 fold change in CNV. Genes called as enhancers (blue) or suppressors (red) are shown in inset table, full results are given in Supplementary Table 3. **(D)** Protein-Protein Physical Interaction Network of factors from the CNV screen and their first neighbours (visualised in Cytoscape Version 3.7.2). Nodes represent proteins and edges represent high-confidence physical interactions between proteins imported from stringApp (v1.6.0). The size and colour of nodes indicates deviation from wild-type fold-change in CNV, with red nodes representing increasing rates of CNV in mutant and blue nodes representing decreasing rates of CNV in mutant. First neighbours of screen factors are shown as small grey nodes. Clusters of interacting proteins (identified in the physical interaction network using ClusterONE v1) which showed similar fold-change in CNV are circled. **(E)** Methodology for assessing CNV and copper adaptation in cells containing 3 copies of the *CUP1* gene (*3xCUP1*). Saturated cultures of *3xCUP1* wild-type and mutant cells were diluted 1:1000 in media ± 0.3 mM CuSO_4_ and incubated for 1 week at 30°C. Half the culture was used for Southern blot analysis, while the other half was diluted 1:80 and grown in a 96-well plate containing a range of concentrations of CuSO_4_ to assay for copper tolerance. **(F)** Southern blot analysis of *CUP1* locus and copper tolerance assay in wild type (wt) and indicated mutant strains with 3 copies of the *CUP1* gene, as outlined in (**E**); Southern blot quantification shows the percentage of alleles deviating from the parental copy number, for adaptation OD_600_ was plotted against [CuSO_4_] and copper tolerance quantified as area-under-curve for each culture; n = 4, except for *pol32Δ* cells where n = 3, p-values calculated by one-way ANOVA.

Seven deletion mutants exhibited a >2-fold increase in CNV, including four DNA replication mutants (*mrc1*Δ, *pol32Δ, sic1Δ* and *rad27Δ)* and three histone deacetylase mutants (*hda1Δ, hda2Δ* and *hst3Δ hst4Δ*). 41 mutants suppressed CNV >2-fold, representing a wider range of biological processes including regulation of transcription (*thp1Δ, med1Δ, cdc73Δ* etc.), DNA repair (*rtt107Δ, rad59Δ, rad52Δ* etc.), nucleosome assembly (*rtt106Δ, cac2Δ, hir1Δ* etc.) and various histone modifiers (*bre1Δ, rtt109Δ, rpd3Δ* etc.) (Figure 1C). ClusterONE, which identifies clusters of interacting proteins in a physical interaction network, revealed clusters of proteins that have similar effects on CNV. Clusters of proteins that promote CNV are involved in chromatin/transcription (orange circle) and DNA repair (purple circle), while a cluster of proteins that facilitate DNA replication & cell cycle progression all tend to suppress CNV (green circle) (Figure 1D). These clusters are coherent with the transcriptionally-stimulated BIR mechanism we have previously proposed (34).

To validate the importance of these genes in adaptive *CUP1* amplification, we introduced individual deletions into a tester strain with 3 copies of *CUP1* (*3xCUP1*) (Figure 1E). During growth in sub-lethal copper, *3xCUP1* cells that undergo *CUP1* amplification gain a selective advantage and become dominant in the population. *CUP1* amplification allows growth in higher concentrations of copper, so cultures grown in sub-lethal copper acquire increased resistance as cells with amplified alleles proliferate. These phenotypes can be quantified by Southern blot and copper resistance assays respectively (Figure 1F).

For example, we included two control strains in the genetic screen that enhance or suppress contraction of 13-copy *CUP1: pol32Δ* and *rtt109Δ* (34). Both alter the frequency at which BIR forks stall and initiate template switching: Pol32 aids the processivity of BIR forks and suppresses template switching (52), while the H3K56 acetylation deposited by Rtt109 promotes template switching (53). If, as we have previously suggested, *CUP1* CNV results from BIR forks encountering H3K56ac then artificially reducing the processivity of BIR forks by deleting *pol32Δ* should promote template switching even in the absence of H3K56ac, and so overcome the suppression of CNV in *rtt109Δ* mutants that lack H3K56ac. We observe exactly this effect in the *3xCUP1* system: deletion of *POL32* rescues both CNV and copper adaptability in *rtt109Δ* (Figure 1F compare *rtt109Δ* to *rtt109Δ pol32Δ),* even though the absence of Pol32 mildly reduces fitness in copper (Figure 1F compare adaptation in wt to *pol32Δ).* This shows that CNV is not impossible in the absence of Rtt109 but rather that *CUP1* CNV occurs through destabilisation of BIR forks by H3K56ac chromatin.

Therefore, our candidate genetic screen for enhancers and suppressors of *CUP1* CNV provides a genetic profile consistent with a transcriptionally stimulated BIR mechanism. Adaptive *CUP1* amplifications are dependent on H3K56ac, which promotes template switching during BIR, but this epigenetic mark is dispensable if the BIR fork is destabilised by loss of Pol32.

### TREX-2 and Mediator are required for transcriptionally stimulated CNV

Many recent studies have shown that RNA:DNA hybrids called R-loops impair replication fork progression, providing a well-validated mechanism for transcription-associated recombination ((54), reviewed in (55)). In consequence, we expected mutants affecting R-loop formation or processing to show strong phenotypes in the genetic screen, but *hpr1Δ, tho2Δ* and *thp2Δ* cells (THO Complex mutants) which accumulate R-loops had little effect (56–58) (Supplementary Table 3). Furthermore, *3xCUP1 rnh1Δ rnh201Δ* cells known to accumulate high levels of R-loops underwent copper adaptation and *CUP1* CNV at normal rates (Figure 2A) (59–61). We therefore find no evidence that R-loops are involved in transcriptionally stimulated *CUP1* CNV.

**Figure 2.**
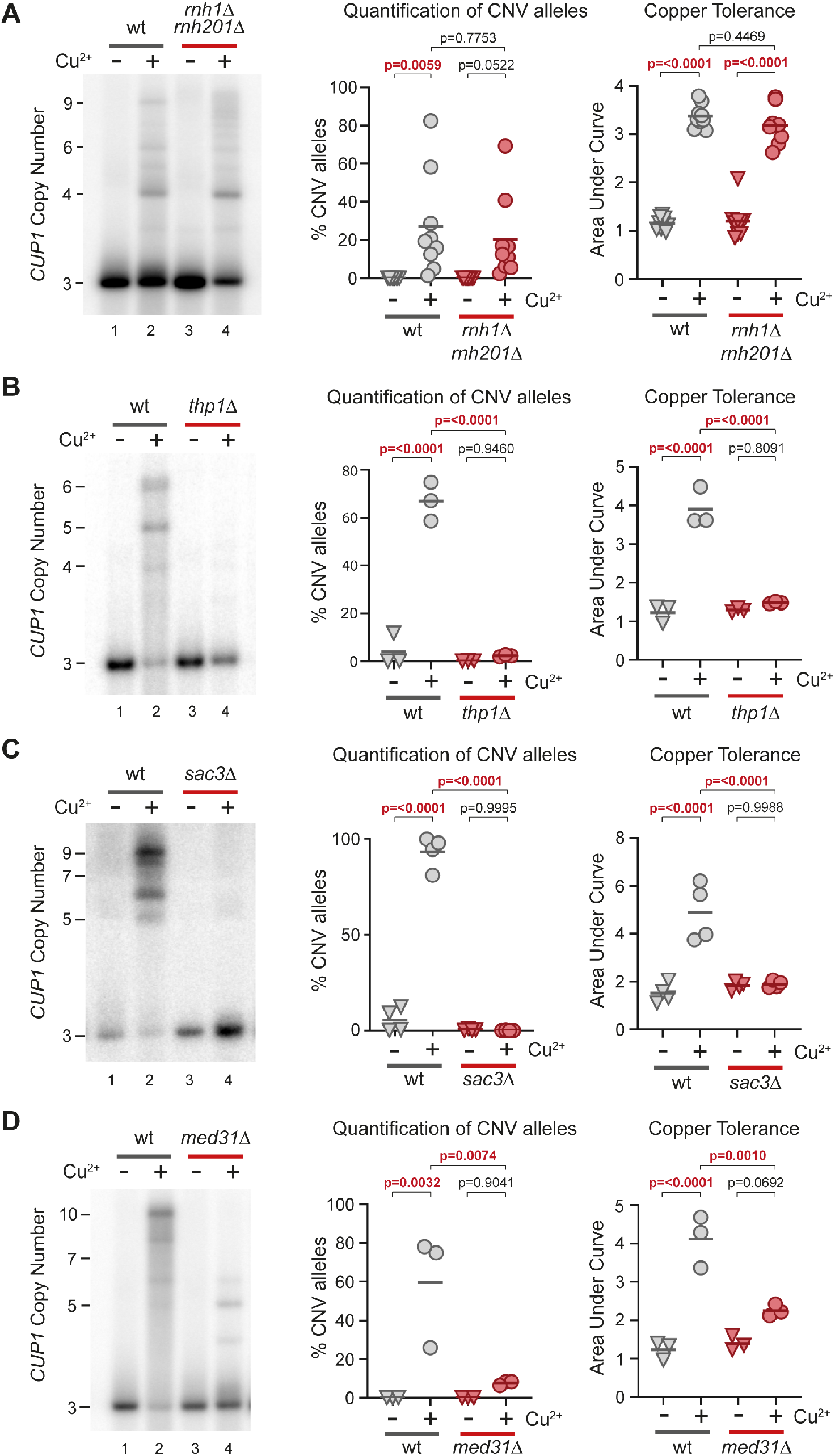
TREX-2 and Mediator are required for transcription stimulated *CUP1* CNV. **(A)** Southern blot analysis of *CUP1* copy number in wild-type (wt) and *rnh1Δ rnh201Δ* cells with 3 copies of the *CUP1* gene, grown to saturation in YNB media ± 0.3 mM CuSO_4_; n = 9. Quantification shows the percentage of alleles deviating from the parental copy number. Copper adaptation was assessed by diluting these same cells in 96-well plates with varying concentrations of CuSO_4_ and incubating for 3 days (see Figure 1E). Final OD_600_ was plotted against [CuSO_4_] and copper tolerance quantified as area-under-curve for each culture; all p-values were calculated by 1-way ANOVA. **(B)** Southern analysis of *CUP1* copy number in *3xCUP1* wild-type (wt) and *thp1Δ* cells after 10 generations ± 0.3 mM CuSO_4_ (Quantification of CNV alleles and Copper adaptation as in **A**); n = 3. **(C)** Southern analysis of *CUP1* copy number in *3xCUP1* wild-type (wt) and *sac3Δ* cells after 10 generations ± 0.3 mM CuSO_4_ (Quantification of CNV alleles and Copper adaptation as in **A**); n = 4. **(D)** Southern analysis of *CUP1* copy number in *3xCUP1* wild-type (wt) and *med31Δ* cells after 10 generations ± 0.3 mM CuSO_4_ (Quantification of CNV alleles and Copper adaptation as in **A**); n = 3.

In contrast, two of the strongest suppressor mutants found in the screen were deletions of *SAC3* and *THP1,* which encode components of Transcription and RNA Export complex 2 (TREX-2) (62,63). Remarkably, *3xCUP1 thp1Δ* and *3xCUP1 sac3Δ* mutants underwent no detectable *CUP1* CNV and were completely unable to adapt to copper despite showing normal resistance to sub-lethal concentrations of copper (Figure 2B/C and Supplementary Figure S2A). Although TREX-2 mutations alter expression of some genes, we did not detect any difference in the induction of *CUP1* in response to copper (Supplementary Figure S2B) (64,65). This implicates TREX-2 as a link between transcription and recombination events at *CUP1.*

TREX-2 is physically associated with the Mediator complex (64) and mediator mutant *med1Δ* also suppressed *CUP1* CNV in the genetic screen. Med1 is a component of the middle module of Mediator but has not been placed within the structure (reviewed in (66)), whereas the TREX-2 / Mediator interface has been mapped to Med31, which projects out from the middle module (64,67,68). Just as we observed in TREX-2 mutants, *3xCUP1 med31Δ* cells did not undergo detectable CNV or adaptation during growth in sub-lethal copper, despite normal *CUP1* mRNA induction (Figure 2D and Supplementary Figure S2B). We further analysed cells lacking Srb2, a component of the head module located at the interface between Mediator and RNA polymerase II (69,70), and found that *3xCUP1 srb2Δ* cells were similarly impaired both in *CUP1* amplification and acquisition of increased copper resistance (Figure S2C). These results reveal a critical role for both TREX-2 and Mediator in the mechanism of *CUP1* amplification.

Mediator connects transcription factors to the core RNA polymerase II machinery at active promoters (reviewed in (71)), while TREX-2 associates with nuclear pores to promote mRNA processing and export (72). However, the gene expression and RNA export functions of Mediator and TREX-2 are separable (64), so the requirement for both complexes in *CUP1* amplification implicates their shared physical interaction. This interaction connects actively transcribed genes to the nuclear pore in a process termed gene gating (64,73–75). Gene gating can impair DNA replication by constraining DNA topology, causing replication forks rendered already fragile by HU treatment to collapse in a Rad53 mutant lacking checkpoint activity (41). This process, albeit only previously characterised under considerable replicative stress, provides a mechanism by which *CUP1* transcriptional induction acting through TREX-2 and Mediator could impede replication forks without a requirement for R-loop formation.

### Homologous recombination occurs at CUP1 without extensive fork stalling or cleavage

We recently developed TrAEL-seq to detect replication fork stalling and replication intermediates (49). Unfortunately, TrAEL-seq is ineffective on copper-treated cells as copper-induced apoptotic fragments generate a high background (49,76) (example data is deposited at GEO GSE154811). However, we have also created a *P_GAL1_-HA cup1* strain in which *GAL1* promoters replace all *CUP1* promoters, and *CUP1* ORFs are replaced by *3HA* ORFs, allowing transcriptional induction at the *cup1* locus using the non-toxic sugar galactose (Figure 3A) (34).

**Figure 3.**
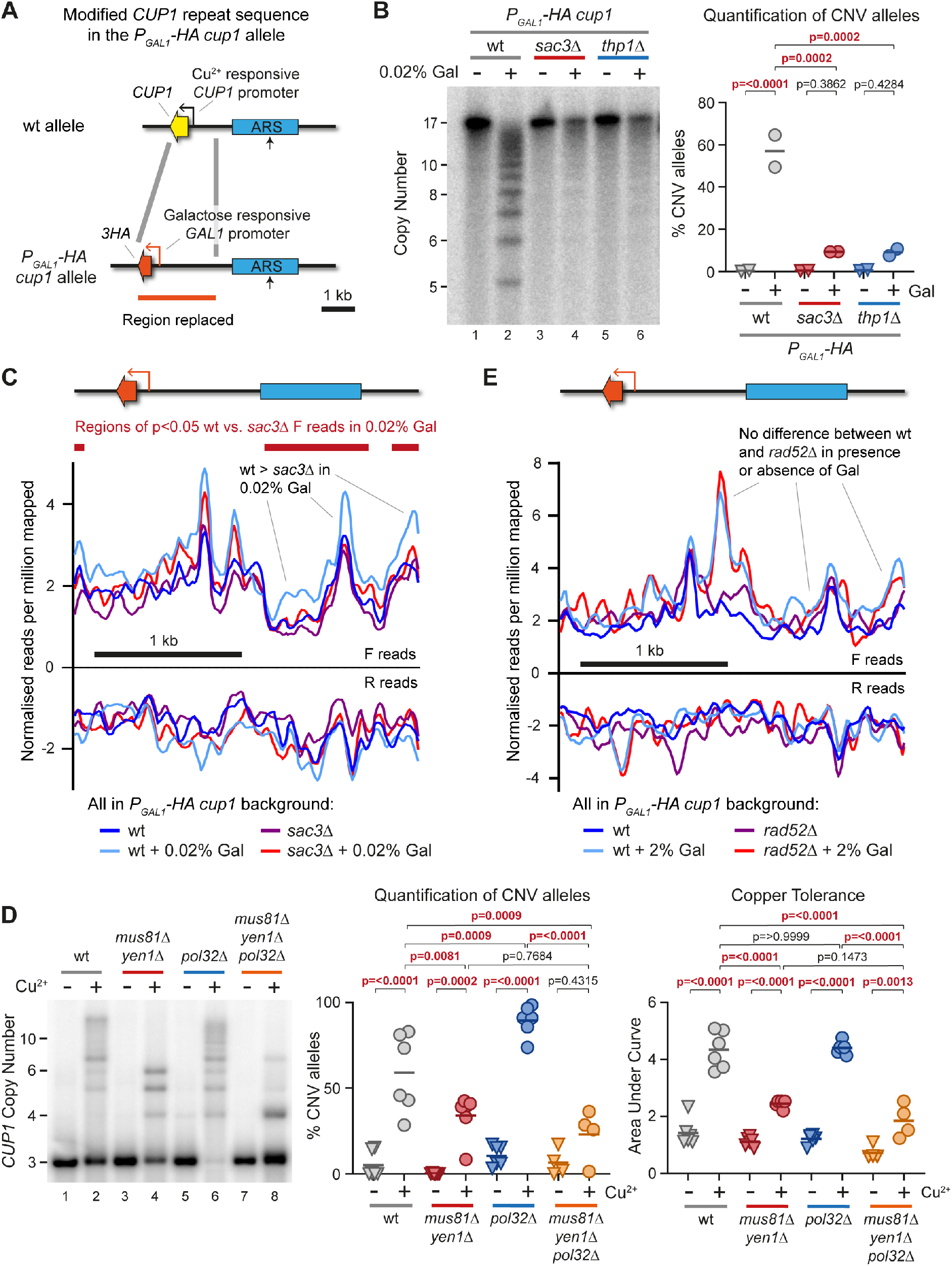
Replication fork stalling and cleavage at *CUP1* locus. **(A)** Schematic of modified *CUP1* repeat in *P_GAL1_-HA cup1* cells, where every *CUP1* ORF is replaced by a 3HA coding sequence (which is phenotypically neutral) and every *CUP1* promoter is replaced by the galactose inducible *GAL1* promoter. **(B)** Southern blot analysis of *P_GAL1_-HA cup1* cells comparing *sac3Δ* and *thp1Δ* cells to wild-type cells grown for 10 generations in 2% raffinose ± 0.02% galactose. Quantification shows the percentage of alleles deviating from the parental copy number of 17 copies; p-values were calculated by one-way ANOVA, n = 2. **(C)** Plots of TrAEL-seq read density on forward and reverse strands across a *P_GAL1_-HA cup1* repeat. Forward TrAEL-seq reads in replicating cells arise primarily from replication forks moving right-to-left, reverse reads from forks moving left-to-right. The dominance of forward over reverse reads shows that replication direction is primarily right-to-left, accumulations of forward reads without a decrease in reverse reads most likely represents increased average fork residency indicative of slower fork progression or more frequent stalling, see Kara *et al.* for more details (49). TrAEL-seq profiles are an average of two biological replicates of *P_GAL1_-HA cup1* wild-type and *sac3Δ* cells grown to mid-log in 2% raffinose, with or without a 6 hour 0.02% galactose induction. Reads were quantified per million reads mapped in 50 bp windows spaced every 10 bp, and an enrichment normalisation applied to make overall read count distributions as uniform as possible, see Figure S3C for the distribution of reads across single copy regions of chromosome VIII. Regions of significant difference between wild type and *sac3Δ* in 0.02% galactose were called using edgeR (50), with forward reads quantified in 200 bp windows spaced every 50 bp, no normalisation was applied prior to the Edge algorithm. **(D)** Southern analysis of *CUP1* copy number in *3xCUP1* wild-type (wt), *mus81Δ yen1Δ, pol32Δ* and *mus81Δ yen1Δ pol32Δ* cells after 10 generations ± 0.3 mM CuSO_4_. Quantification shows the percentage of alleles deviating from the parental copy number, p-values calculated by 1-way ANOVA. Copper adaptation was assessed by treated cells with varying concentrations of CuSO_4_ and incubating for 3 days. Final OD_600_ was plotted against [CuSO_4_] and copper tolerance quantified as area-under-curve for each culture; p-values were calculated by one-way ANOVA; n = 4. **(E)** Plots of TrAEL-seq read density on forward and reverse strands across a *P_GAL1_-HA cup1* repeat as in **C**, for wild-type or *rad52Δ* cells grown to mid log in 2% raffinose followed by either 2% glucose or 2% galactose for 6 hours, n=1.

TrAEL-seq reads accumulate at sites of replication fork stalling, including at the endogenous *GAL1* promoter when transcriptionally active, providing a measure of disruption to replication fork progression (49). *GAL1* promoter activity is exceptionally strong under normal 2% galactose induction, leading to saturating levels of Sac3- and Thp1-independent CNV in 10 generations that probably result from direct collisions between RNA polymerase and the replication fork (Supplementary Figure S3A). However, CNV is Sac3- and Thp1-dependent under moderate 0.02% galactose induction that should better reflect *CUP1* gene expression (Figure 3B), so we constructed TrAEL-seq libraries from *P_GAL1_-HA cup1* wild-type and *sac3Δ* cells induced with 0.02% galactose.

We expected to observe a pronounced accumulation of TrAEL-seq reads at the *P_GAL1_-HA* promoter as a result of replication fork stalling, but the promoter peaks formed in 0.02% galactose were modest and Sac3-independent (Figure 3C), although further enhanced in 2% galactose (Supplementary Figure S3B). In contrast, the region upstream of the *P_GAL1_* promoter containing the ARS element accumulated significantly more TrAEL-seq reads in wild type than *sac3Δ* cells on induction with 0.02% galactose (right hand side and far left of Figure 3C). Normalisation errors could also give rise to such a baseline signal increase, however global TrAEL-seq read distributions were unaffected by addition of galactose (Supplementary Figure S3C shows all equivalent windows in single copy regions of chromosome VIII), and if anything the normalisation process globally increases *sac3Δ* signals compared to wild-type. Therefore, we observe a small Sac3-dependent increase in TrAEL-seq read density outside the transcribed region and particularly around the ARS element in *P_GAL1_-HA cup1* that is consistent with replication forks stalling but not at well-defined sites.

Homologous recombination can be initiated from stalled forks, and both *CUP1* amplification and acquisition of copper resistance are strongly dependent on the homologous recombination proteins Rad52 and Rad51 that mediate strand invasion (Supplementary Figure S3D). A one-ended DSB for strand invasion can be formed through cleavage of the stalled replication fork by structure specific endonucleases (SSEs) Mus81, Yen1 or Slx1/Slx4 (reviewed in (77)). Deletion of *MUS81* alone had little effect on *CUP1* CNV, but a double deletion of *MUS81* and *YEN1* strongly suppressed CNV and adaptation (Figure 3D, Supplementary Figure S3E). Unlike Rtt109, which has a regulatory role in CNV, deletion of *POL32* to drive constitutive template switching did not restore CNV in cells lacking Mus81 and Yen1, showing that SSE activity is important for the CNV mechanism (Figure 3D). If SSEs cleave stalled replication forks to initiate homologous recombination then cleaved DNA ends should accumulate in a *rad52*Δ mutant that cannot undergo strand invasion. However, we detected no additional accumulation of TrAEL-seq reads in *P_GAL1_-HA cup1 rad52Δ* cells compared to wild type even under 2% galactose induction which drives extremely high levels of CNV (Figure 3E). This indicates that SSEs act after the initiation of homologous recombination rather than at the initiation step.

The Sac3-dependent increase in TrAEL-seq reads across *Pgal-HA cup1* is consistent with replication forks stalling due to DNA topological constraint, which would not happen at a defined location but rather at random across a wide area. The dependence of *CUP1* amplification on homologous recombination and SSEs indicates that BIR is induced from stalled forks; however, we do not detect fork cleavage intermediates in a *rad52Δ* mutant. While the absence of TrAEL-seq read accumulation does not prove that fork cleavage is absent, it does suggest that other mechanisms are more important for generating the very high levels of CNV observed under galactose induction (Figure 3B). BIR does not necessarily require fork cleavage (23) and our observations are more coherent with Mus81 and Yen1 acing in resolution rather than initiation of recombination (78,79).

### *Fork progression and replication timing are critical mediators of* CUP1 *CNV*

Gene gating is widespread (65) and if gene gating alone invoked the levels of recombination we observe at *CUP1* then dramatic genomic instability would ensue. We therefore examined replication at *CUP1* to determine whether locus specific factors are also involved in transcriptionally stimulated CNV, focusing on replisome component Mrc1 as the strongest negative regulator of *CUP1* CNV in the genetic screen.

Mrc1 mediates the DNA replication checkpoint, which modulates the DNA replication programme in response to replication fork stalling (80,81). Mec1 phosphorylates Mrc1 in response to single stranded DNA accumulation, and then phosphorylated Mrc1 binds to and activates Rad53 (reviewed in (82)). We therefore assayed copper adaptation in *rad53Δ* and *mec1Δ* mutants using a *3xCUP1 sml1Δ* background as these mutations are lethal without *SML1* deletion (83) (Supplementary Figure S4A). Loss of Mec1 reduced CNV without affecting adaptation indicating a positive but non-essential role for DNA damage signalling in CNV, whilst *rad53Δ* had no effect on CNV or adaptation, which is not entirely surprising as Rad53 activation requires a multiple stalled replication forks (84). The mildly reduced *CUP1* CNV phenotype of *mec1Δ* is in stark contrast to the dramatic acceleration of CNV and adaptation in *3xCUP1 mrc1Δ* cells (Figure 4A), so the checkpoint mediator function of Mrc1 cannot explain the CNV phenotype of *mrc1Δ.*

**Figure 4.**
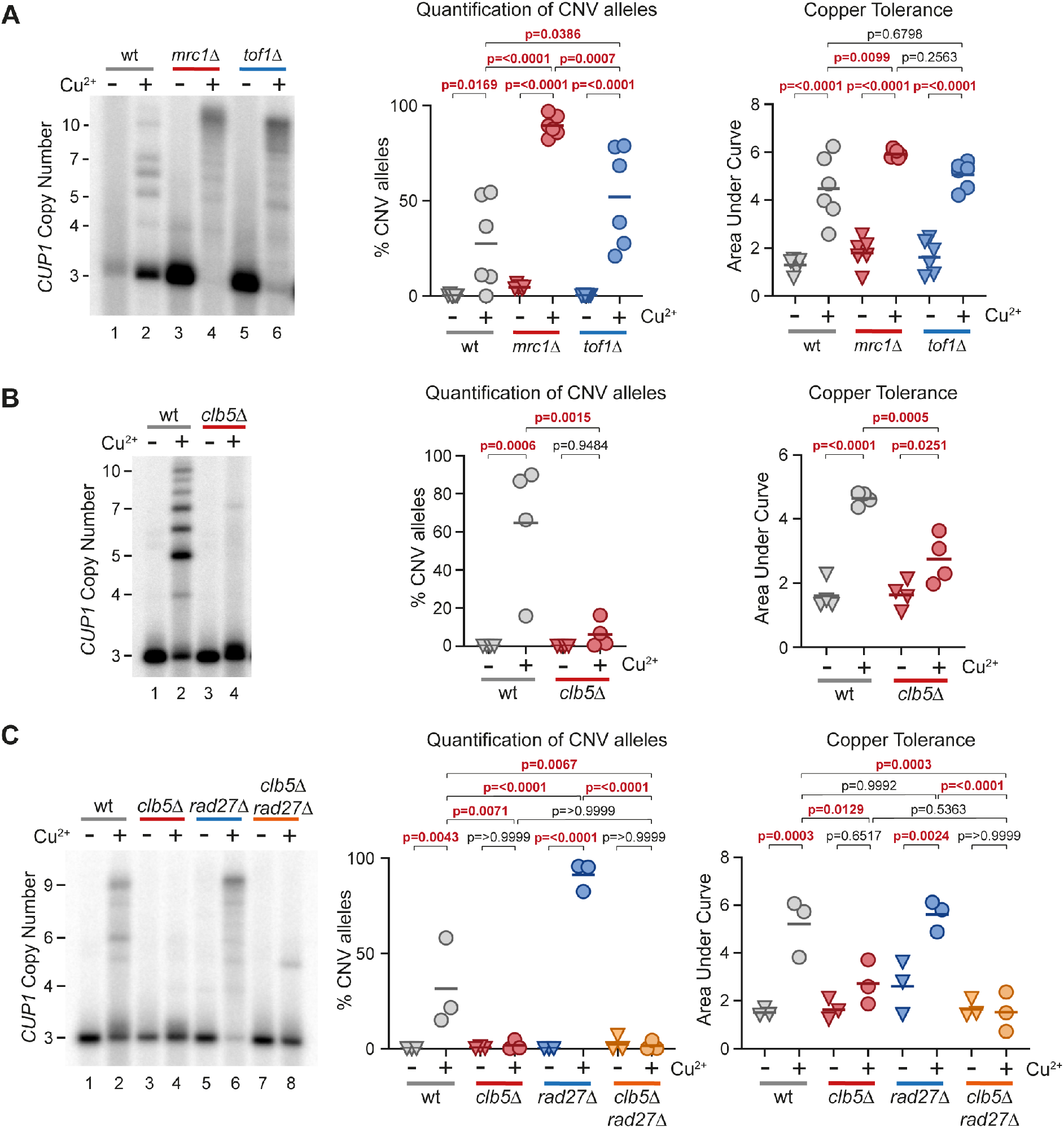
Effect of replication timing and replication fork progression on *CUP1* CNV. **(A)** Southern analysis of *CUP1* copy number in *3xCUP1* wild-type (wt), *mrc1Δ* and *tof1Δ* cells after 10 generations ± 0.3 mM CuSO_4_. Quantification shows the percentage of alleles deviating from the parental copy number, p-values calculated by 1-way ANOVA. Copper adaptation was assessed by treating cells with varying concentrations of CuSO_4_ for 3 days, final OD_600_ was plotted against [CuSO_4_] and copper tolerance quantified as area-under-curve for each culture; p-values were also calculated by one-way ANOVA; n = 6. **(B)** Southern blot analysis of *CUP1* copy number and growth curve analysis in *3xCUP1* wild type (wt) and *clb5Δ* mutants. Cells were cultured and CNV alleles and copper adaptation quantified as in **A**; n = 4. **(C)** Southern and growth curve analysis of *3xCUP1* wild type (wt) and indicated mutant cells. Cells were cultured, CNV alleles and copper adaptation quantified as in **A**; n = 3.

Mrc1 is also important for replication fork progression: Mrc1, Tof1 and Csm3 form the Fork Protection Complex and together regulate replication fork speed (85,86), whilst Mrc1 and Tof1 have important but differing roles during replication fork stalling. Tof1 stabilises stalled forks at programmed pause sites, whereas Mrc1 stabilises the replisome during unscheduled fork stalling (87,88). Deletion of *TOF1* in *3xCUP1* significantly accelerated *CUP1* CNV similarly to *mrc1Δ,* which suggests that replication fork speed rather than stabilisation of a particular class of stalled fork is the primary determinant of CNV rate (Figure 4A). A reduction in replication fork speed would increase the chance that late replicating regions are not replicated prior to G2/M, requiring emergency repair by BIR as at mammalian fragile sites (89). The *CUP1* locus is in a particularly late replicating region of Chromosome VIII (Supplementary Figure S4B), which provided a possible explanation for the high frequency of CNV events.

If CNV arises through late replication then mutations that extend S-phase should suppress *CUP1* CNV by increasing the time available for replication. Clb5 is a cyclin that promotes activation of late firing replication origins towards the end of S phase; replication fork progression is normal in *clb5Δ* mutants (90) but loss of Clb5 prevents firing of late origins and S phase is extended to allow completion of replication (91,92). Remarkably, *CUP1* CNV decreased substantially in *clb5Δ* cells with a concurrent reduction in copper adaptation (Figure 4B), showing that Clb5 activity is important for *CUP1* CNV. To confirm this result we tested two other mutants, *sic1Δ* and *dia2Δ,* which cause similar shifts in cell cycle profile to *clb5Δ* (93) albeit through a variety of mechanisms, and obtained the same result (Supplementary Figure S4C). This suggested that S-phase duration controls *CUP1* CNV rate.

If *clb5Δ* extends S-phase but *mrc1Δ* slows fork progression, we predicted that the *CUP1* CNV phenotype of an *mrc1Δ clb5Δ* double mutant would approach wild type. Unfortunately, this double mutant was not viable in the *3xCUP1* background but we achieved a similar outcome by combining *clb5Δ* with *rad27Δ,* another replisome mutation that dramatically enhanced *CUP1* CNV in the genetic screen. Rad27 is the budding yeast FEN1 ortholog required for efficient processing of Okazaki fragments at replication forks. Loss of Rad27 slows fork progression (93,94), increases replication stress, and impairs DNA repair leading to high levels of genome instability through multiple mechanisms (95,96), so the high rate of *CUP1* CNV was at first sight unsurprising (Figure 4C). However, the combination of *clb5Δ* with *rad27Δ* completely suppressed *CUP1* CNV and copper adaptation (Figure 4C), showing that acceleration of CNV in *rad27Δ* occurs through the same mechanism as CNV in wild-type cells rather than unrelated defects in Okazaki fragment synthesis or DNA repair. Nonetheless, this result was very puzzling as it is hard to reconcile the complete suppression of CNV in *clb5Δ rad27Δ* with a model in which Clb5 simply extends replication timing to allow replication forks more time to finish DNA synthesis; we would not expect this to be sufficient to offset the loss of Rad27.

These observations show that *CUP1* CNV rates are very sensitive to normal replication fork progression, with slowing of replication forks promoting rapid CNV and copper adaptation. The efficient repression of *CUP1* CNV by mutants that affect origin activity and S-phase duration shows that encounters between normal replication forks and highly transcribed loci do not cause CNV in isolation. Rather, subtle changes to the replication programme are sufficient to suppress the CNV mechanism meaning that homologous recombination must only occur under very specific circumstances.

### Late firing replication origins rather than late replication are the primary driver of CUP1 CNV

The unexpectedly strong suppression of CNV in *clb5Δ rad27Δ* led us to examine the replication profile of the *CUP1* locus more closely. Extension of S-phase in *clb5Δ* and *sic1Δ* mutants is attributed to suppression of late-firing replication origins (92,93,97), and conversely deletion of *MRC1* increases usage of late firing origins (81,93). It is therefore possible that usage of late firing local origins is the principle determinant of *CUP1* CNV, rather than replication timing or fork speed. Importantly, despite the late replication of the locus, replication origins (ARS elements) are present in each of the *CUP1* repeats although whether these fire during normal replication has remained unclear (Figure 1A).

Formation of a replication bubble at a replication origin creates two replication forks that move in opposite directions, and replication origins are therefore detectable as sites of sharp changes in replication fork direction. Replication fork direction is revealed by the polarity of TrAEL-seq reads (49), and TrAEL-seq data from BY4741 wild-type cells with 13 copies of *CUP1* shows a change in polarity from negative to positive at the *CUP1* origin indicative of ARS activity (Figure 5A upper panel, highlighted in orange). This signal is weak compared to other ARS elements in the region and becomes undetectable in *3xCUP1* cells (Supplementary Figure 5A), consistent with the *CUP1* ARS elements being functional but rarely active. In contrast, very little change in replication fork polarity is detected at *CUP1* in *clb5Δ* cells with 13 copies of *CUP1,* placing the *CUP1* ARS elements amongst those late firing origins dependent on Clb5 (Figure 5A lower panel). We note that the two closest origins to *CUP1* were also repressed in *clb5Δ* (Figure 5A, at 133 and 286 kb), so although loss of Clb5 extends S-phase it is questionable whether *CUP1* replicates any earlier in this mutant as the replication forks which replicate this region must travel further (from ARS elements at ~64 and 392 kb).

**Figure 5.**
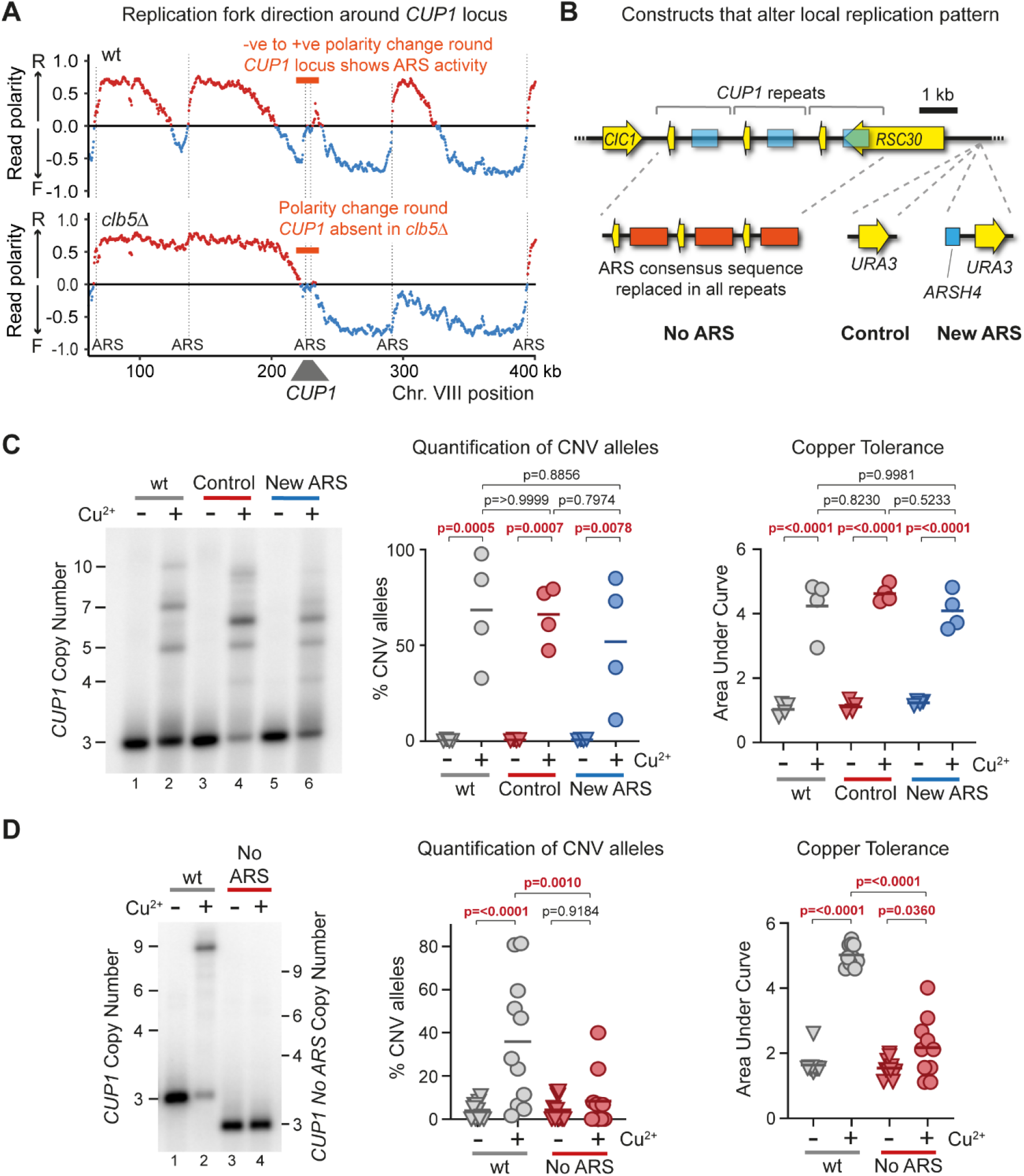
Local Replication origin firing regulates CNV at *CUP1* locus. **(A)** TrAEL-seq read polarity plots for wild type and *clb5Δ* cells grown to mid log in 2% Glucose. TrAEL-seq detects replication direction, an excess of reverse (R) reads at any site (indicated by positive polarity, red), results from replication forks moving left-to-right, the opposite for replication forks moving right-to-left (blue). Sharp transitions from −ve to +ve occur at active replication origins (ARS elements), gradual transitions from +ve to −ve are regions where forks converge. Plots show regions surrounding *CUP1* on chromosome VIII, quantified by (R-F)/(R+F) with R and F referring to Reverse and Forward reads respectively; n = 2, data from GSE154811. **(B)** Schematic of the 3 copy *CUP1* array and the modifications introduced to influence local replication pattern. In “No ARS” cells, the region containing the endogenous ARS upstream of every *CUP1* repeat have been replaced by a non-expressed sequence derived from a *GFP* tagging plasmid. “New ARS” and “Control” cells have a *URA3* marker integrated upstream of the *RSC30* promoter, either with or without an efficient replication origin (*ARSH4)* respectively. TrAEL-seq data confirming activity of the new ARS is shown in Figure S5A. **(C)** Southern blot analysis of *CUP1* copy number in *3xCUP1* wild-type (wt), Control and New ARS cells (described in **B**). Cells were cultured for 10 generations ± 0.3 mM CuSO_4_. Quantification shows the percentage of alleles deviating from the parental copy number, p-values calculated by 1-way ANOVA. Copper adaptation was assessed by treating cells with varying concentrations of CuSO_4_ and incubating for 3 days, final OD_600_ was plotted against [CuSO_4_] and copper tolerance quantified as area-under-curve for each culture; p-values were also calculated by 1-way ANOVA; n = 4. **(D)** Southern blot analysis of *CUP1* copy number and growth curve analysis in *3xCUP1* wild-type (wt) and *3xCUP1* cells with the region containing the ARS sites replaced by unrelated sequence (described in **B**). Cells were cultured, CNV alleles and copper adaptation quantified as in **B**, n = 10 for Southern blot analysis, n = 11 for Growth curve analysis.

The presence of Clb5-dependent ARS elements at *CUP1* is consistent with the possibility that *CUP1* ARS activity is important for *CUP1* CNV. If so then loss of Clb5 would suppress CNV by preventing local ARS activity rather than by extending S phase. To distinguish these possibilities, we created origin insertion and deletion strains that either prevent *CUP1* ARS activity or force early replication (Figure 5B).

Firstly, to promote early replication of *CUP1* we integrated the efficient and early *ARSH4* origin adjacent to the *CUP1* locus to create a “New ARS” strain, along with a control containing the selectable marker but no new ARS element (Figure 5B). TrAEL-seq analysis confirmed that this origin was efficiently activated and dominated the local replication profile (Supplementary Figure S5A compare middle and bottom panels). However, this had no detectable impact on copper adaptation or *CUP1* repeat amplification (Figure 5C). Therefore, *CUP1* amplification does not depend on replication forks arriving at the locus late in S-phase from distant origins, and the extension of S-phase in *clb5Δ* is unlikely to explain the strong suppression of CNV.

Secondly, to prevent replication origin activity in *CUP1* we mutated the ARS element in each *CUP1* repeat. As there are considerable discrepancies in the locations assigned to the *CUP1* ARS (98,99) we opted to delete a substantial region of each *CUP1* repeat upstream of the *CUP1* promoter, replacing this with unrelated sequence derived from a *GFP* construct (Figure 5B “No ARS”). This change did not alter basal copper resistance compared to *3xCUP1* cells containing wild type *CUP1* repeats (Supplementary Figure 5B top panel), showing that *CUP1* function was unaffected. However, when pre-exposed to copper, loss of the ARS region suppressed *CUP1* CNV and copper adaptation similarly to *CLB5* deletion (Figure 5D). This shows that activity of local ARS elements in the *CUP1* repeats, which is suppressed in *clb5Δ,* drives *CUP1* CNV.

These experiments resolve the unexpected outcomes of *CLB5* deletion and show that firing of inefficient replication origins at *CUP1* rather than late replication timing underlies the replication-dependence of transcriptionally stimulated *CUP1* amplification. Taken together with our previous results, this suggests that replication origin firing, in regions of high topological strain caused by promoter-nuclear pore interactions, promotes template switching events that result in adaptive CNV.

## Discussion

Here we have dissected the mechanism by which transcriptional induction of the copper resistance gene *CUP1* stimulates CNV events that cause *CUP1* amplification and thereby increase copper resistance. We demonstrate critical roles for the TREX-2 and Mediator complexes that link transcribed loci to the nuclear pore, and for local replication origin activity, in addition to the known importance of H3K56ac.

### A two-step mechanism for CUP1 CNV

Given recent discoveries of replication fork instability resulting from collisions between replication forks and RNA polymerase II or R-loops, the results of our genetic screen for *CUP1* CNV modulators were not what we expected (29,30,39,40,43). The importance of TREX-2 and Mediator, deletions of which are even stronger suppressors of copper adaptation than *rad52Δ,* suggests that topological constraint of replication fork progression causes CNV, rather than direct interactions with the transcription unit, and this is coherent with the small but widespread increase in TrAEL-seq read density when the locus is transcriptionally active. Nonetheless, topological impairment of replication fork progression is known (41,42) and it is not is not too surprising that this increases the frequency of homologous recombination events that can result in CNV.

However, the requirement for H3K56ac needs to be explained (34). H3K56ac promotes template switching during BIR (53), but given that H3K56ac is present on new histones that are deposited behind the replication fork we must explain how a migrating D-loop formed during fork repair, which should use unreplicated DNA ahead of the stalled fork as a template, encounters H3K56ac chromatin. In other words, how is H3K56ac deposited ahead of the replication fork? The activity of local replication origins in regions of frequent replication fork stalling provides one solution to this: if both forks in the nascent replication bubble stall then the replication bubble could dissolve by fork reversal (illustrated in Figure S6). However, during the formation and initial progression of the bubble, histone exchange would leave an “epigenetic scar” of H3K56ac in a non-replicated genomic region (also illustrated in Figure S6). In theory, *CUP1* transcription could also cause histone exchange in front of the fork, but copper resistance is not reduced when the upstream region containing the replication origin is removed whereas copy number amplification is suppressed.

The mechanism we propose for transcriptionally-stimulated *CUP1* CNV is illustrated in Figure 6. Firstly, a replication origin fires in the *CUP1* region but rapidly stalls and the bubble collapses due to replicative stress imposed by TREX-2 and Mediator. Firing of a second origin again leads to inefficient progression, but if this bubble does not collapse then the fork must be restarted by fork reversal and D-loop formation. This D-loop becomes prone to template switching on encountering the epigenetic scar of H3K56ac left by the first collapsed replication bubble, increasing the frequency of local CNV events.

**Figure 6.**
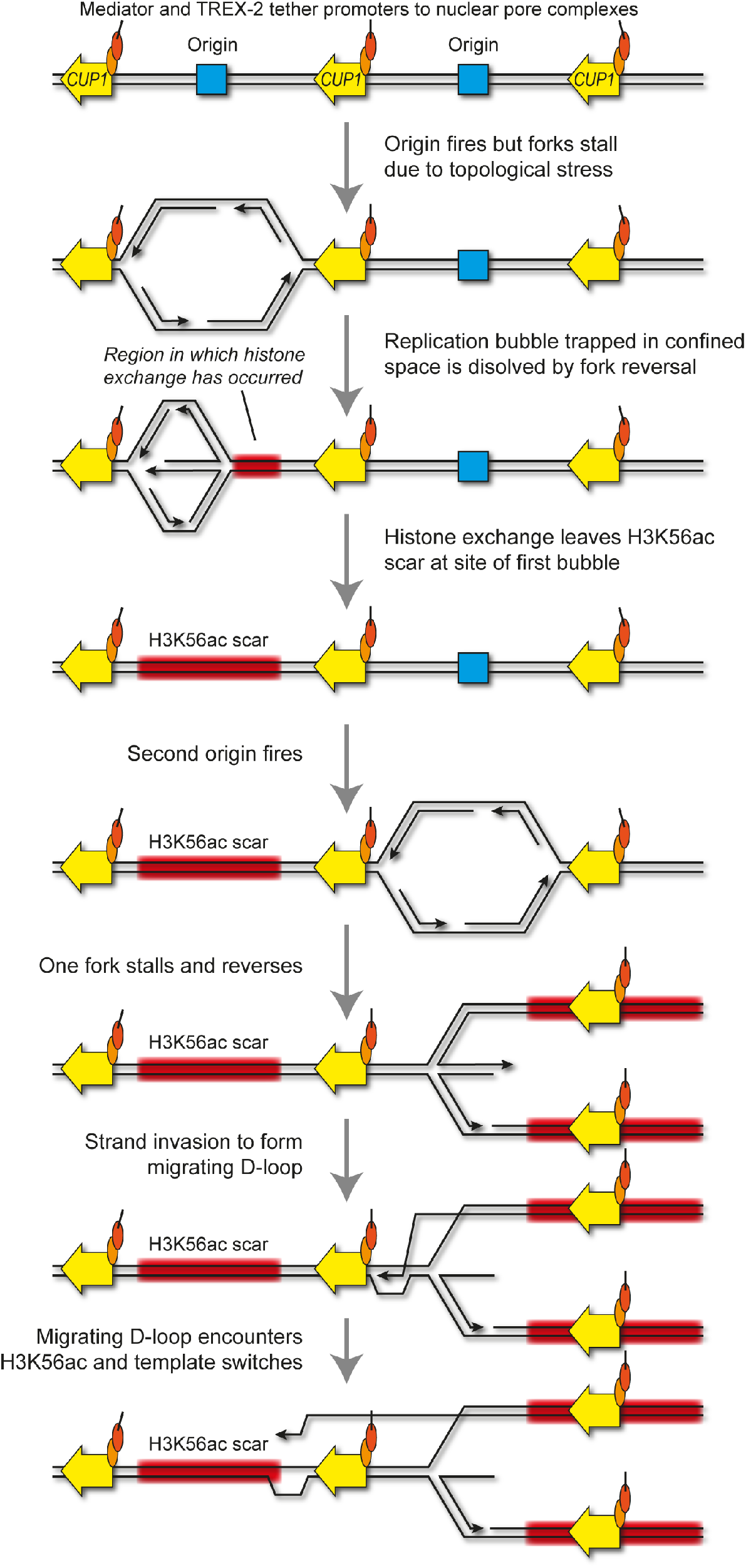
A model for origin-dependant stimulated-CNV of *CUP1* locus. DNA strands are shown in black, the *CUP1* gene shown in yellow, replication origins are shown in blue, chromatin containing H3K56ac shown in red, and mediator and TREX2 complexes are shown as orange and red circles respectively. Tethering of *CUP1* array to the nuclear pore during gene gating may promote topological stresses causing some replication forks to stall and potentially dissolve, leaving behind a H3K56ac scar. Replication forks that encounter H3K56ac chromatin are prone to template switches that can result in non-allelic homologous recombination leading to CNV.

We note that this mechanism contrasts with recent studies showing that nuclear pore localisation enhances DNA repair (100) and suppresses R-loops (101). Indeed, it is clear that more than one mutational pathway acts even at *CUP1;* we focus here on adaptive *CUP1* amplification and find this to be Rad51- and Sic1-dependent, but maladaptive *CUP1* contractions assayed in our screen and another recent study (102) are Rad51-independent, Rad59-dependent and suppressed by Sic1. This provides a sobering insight into the complexity of replication-linked repair events.

### Replication stress as a driver of adaptation

Adaptive mutation at *CUP1* does not fit classical models of adaptation through selection of random pre-existing mutations, as *CUP1* CNV occurs in response to copper stress. However, neither does this constitute a stress-induced mutational pathway equivalent to the bacterial SOS response or recent reports of adaptive mutability during chemotherapy (103–105). Stress induced mutation pathways invoke a switch to the use of mutagenic repair pathways for repair of random DNA damage (reviewed in (106)), whereas the frequency and location of DNA damage is dramatically increased by the replication conflicts with a highly transcribed *CUP1* allele, driving CNV without necessitating a change in repair mechanism.

More generally, we suggest that DNA replication under environmental challenge is inherently mutagenic and prone to increase the genetic heterogeneity of a population from which adaptive mutations can be selected. Cells exposed to imperfect environments may be metabolically suboptimal and have reduced levels of dNTPs, ATP or histones needed for replication, may strongly induce stress responsive genes orientated head-on to replication forks (as in bacteria (107)) and may have increased levels of DNA lesions that impair replication fork progression. These would reduce fork progression and increase the frequency of local repair events, but also incur the widespread usage of inefficient origins that we find to be mutagenic at *CUP1* and increase the likelihood of fragile sites not completing replication at G2/M (89,108–113). Therefore, increased mutation rate does not need to be actively stimulated or initiated by cells under environmental challenge as long as some level of replication is maintained, since increased genetic heterogeneity should be an emergent property of replication under stress.

The question of when and how resistance mutations form is critical; as yeast pathogens become resistant to fungicides we will face major challenges in medicine and food production (reviewed in (114)). An example of the damage wrought by fungal pathogens is the progressive annihilation of amphibians by *Batrachochytrium dendrobatidis* (reviewed in (115)). Here we provide proof of principle that the emergence of adaptive mutations is not inevitable as multiple genetic manipulations prevent budding yeast acquiring resistance to the agricultural fungicide copper sulfate. If adaptive mutations generally arise during fungicidal treatment in medicine or agriculture then a window of opportunity for avoiding resistance exists: by altering or supplementing treatment regimens to prevent replication or diminish replicative stress, the occurrence of *de novo* mutation should be reduced.

## Acknowledgements

We would like to thank Neesha Kara for help with TrAEL-seq and generation of Read Polarity plots, Paula Koko Gonzales and Nicole Forrester of the Babraham Institute Next Generation Sequencing for data generation, and Claudia Ribeiro de Almeida for critical reading of the manuscript.

## Funding

JH, AW, MK and RH were funded by the Wellcome Trust [110216], JH and FK by the BBSRC [BI Epigenetics ISP: BBS/E/B/000C0423]. The funders had no role in study design, data collection and analysis, decision to publish, or preparation of the manuscript.

This research was funded in whole, or in part, by the Wellcome Trust [Grant number 110216]. For the purpose of open access, the author has applied a CC BY public copyright licence to any Author Accepted Manuscript version arising from this submission.

## Notes

### Competing Interest Statement

The authors have declared no competing interest.

## References

1. Katju, V. and Bergthorsson, U. (2013) Copy-number changes in evolution: rates, fitness effects and adaptive significance. Front Genet, 4, 273.

2. Frei, E., 3rd, Rosowsky, A., Wright, J.E., Cucchi, C.A., Lippke, J.A., Ervin, T.J., Jolivet, J. and Haseltine, W.A. (1984) Development of methotrexate resistance in a human squamous cell carcinoma of the head and neck in culture. Proc Natl Acad Sci U S A, 81, 2873–2877.

3. Corcoran, R.B., Dias-Santagata, D., Bergethon, K., Iafrate, A.J., Settleman, J. and Engelman, J.A. (2010) BRAF gene amplification can promote acquired resistance to MEK inhibitors in cancer cells harboring the BRAF V600E mutation. Sci Signal, 3, ra84.

4. Huennekens, F.M. (1994) The methotrexate story: a paradigm for development of cancer chemotherapeutic agents. Adv Enzyme Regul, 34, 397–419.

5. Little, A.S., Balmanno, K., Sale, M.J., Newman, S., Dry, J.R., Hampson, M., Edwards, P.A., Smith, P.D. and Cook, S.J. (2011) Amplification of the driving oncogene, KRAS or BRAF, underpins acquired resistance to MEK1/2 inhibitors in colorectal cancer cells. Sci Signal, 4, ra17.

6. Sandegren, L. and Andersson, D.I. (2009) Bacterial gene amplification: implications for the evolution of antibiotic resistance. Nat Rev Microbiol, 7, 578–588.

7. Selmecki, A., Forche, A. and Berman, J. (2006) Aneuploidy and isochromosome formation in drug-resistant Candida albicans. Science, 313, 367–370.

8. Lucas, E.R., Miles, A., Harding, N.J., Clarkson, C.S., Lawniczak, M.K.N., Kwiatkowski, D.P., Weetman, D., Donnelly, M.J. and Anopheles gambiae Genomes, C. (2019) Whole-genome sequencing reveals high complexity of copy number variation at insecticide resistance loci in malaria mosquitoes. Genome Res, 29, 1250–1261.

9. Jones, L., Riaz, S., Morales-Cruz, A., Amrine, K.C., McGuire, B., Gubler, W.D., Walker, M.A. and Cantu, D. (2014) Adaptive genomic structural variation in the grape powdery mildew pathogen, Erysiphe necator. BMC Genomics, 15, 1081.

10. Zhang, F., Gu, W., Hurles, M.E. and Lupski, J.R. (2009) Copy number variation in human health, disease, and evolution. Annu Rev Genomics Hum Genet, 10, 451–481.

11. Hastings, P.J., Ira, G. and Lupski, J.R. (2009) A microhomology-mediated break-induced replication model for the origin of human copy number variation. PLoS Genet, 5, e1000327.

12. Gu, W., Zhang, F. and Lupski, J.R. (2008) Mechanisms for human genomic rearrangements. Pathogenetics, 1, 4.

13. Liu, P., Carvalho, C.M., Hastings, P.J. and Lupski, J.R. (2012) Mechanisms for recurrent and complex human genomic rearrangements. Curr Opin Genet Dev, 22, 211–220.

14. Hastings, P.J., Lupski, J.R., Rosenberg, S.M. and Ira, G. (2009) Mechanisms of change in gene copy number. Nat Rev Genet, 10, 551–564.

15. Zhang, F., Carvalho, C.M. and Lupski, J.R. (2009) Complex human chromosomal and genomic rearrangements. Trends Genet, 25, 298–307.

16. Mazzagatti, A., Shaikh, N., Bakker, B., Spierings, D.C.J., Wardenaar, R., Maniati, E., Wang, J., Boemo, M.A., Foijer, F. and McClelland, S.E. (2020) DNA Replication Stress Generates Distinctive Landscapes of DNA Copy Number Alterations and Chromosome Scale Losses. bioRxiv.

17. Arlt, M.F., Ozdemir, A.C., Birkeland, S.R., Wilson, T.E. and Glover, T.W. (2011) Hydroxyurea induces de novo copy number variants in human cells. Proc Natl Acad Sci U S A, 108, 17360–17365.

18. Arlt, M.F., Mulle, J.G., Schaibley, V.M., Ragland, R.L., Durkin, S.G., Warren, S.T. and Glover, T.W. (2009) Replication stress induces genome-wide copy number changes in human cells that resemble polymorphic and pathogenic variants. Am J Hum Genet, 84, 339–350.

19. Petermann, E. and Helleday, T. (2010) Pathways of mammalian replication fork restart. Nat Rev Mol Cell Biol, 11, 683–687.

20. Hanada, K., Budzowska, M., Davies, S.L., van Drunen, E., Onizawa, H., Beverloo, H.B., Maas, A., Essers, J., Hickson, I.D. and Kanaar, R. (2007) The structure-specific endonuclease Mus81 contributes to replication restart by generating double-strand DNA breaks. Nat Struct Mol Biol, 14, 1096–1104.

21. Froget, B., Blaisonneau, J., Lambert, S. and Baldacci, G. (2008) Cleavage of stalled forks by fission yeast Mus81/Eme1 in absence of DNA replication checkpoint. Mol Biol Cell, 19, 445–456.

22. Ait Saada, A., Lambert, S.A.E. and Carr, A.M. (2018) Preserving replication fork integrity and competence via the homologous recombination pathway. DNA Repair (Amst), 71, 135–147.

23. Carr, A.M. and Lambert, S. (2013) Replication stress-induced genome instability: the dark side of replication maintenance by homologous recombination. J Mol Biol, 425, 4733–4744.

24. Smith, C.E., Llorente, B. and Symington, L.S. (2007) Template switching during break-induced replication. Nature, 447, 102–105.

25. Lydeard, J.R., Jain, S., Yamaguchi, M. and Haber, J.E. (2007) Break-induced replication and telomerase-independent telomere maintenance require Pol32. Nature, 448, 820–823.

26. Sakofsky, C.J., Roberts, S.A., Malc, E., Mieczkowski, P.A., Resnick, M.A., Gordenin, D.A. and Malkova, A. (2014) Break-induced replication is a source of mutation clusters underlying kataegis. Cell Rep, 7, 1640–1648.

27. Deem, A., Keszthelyi, A., Blackgrove, T., Vayl, A., Coffey, B., Mathur, R., Chabes, A. and Malkova, A. (2011) Break-induced replication is highly inaccurate. PLoS Biol, 9, e1000594.

28. Saini, N., Ramakrishnan, S., Elango, R., Ayyar, S., Zhang, Y., Deem, A., Ira, G., Haber, J.E., Lobachev, K.S. and Malkova, A. (2013) Migrating bubble during break-induced replication drives conservative DNA synthesis. Nature, 502, 389–392.

29. Lang, K.S., Hall, A.N., Merrikh, C.N., Ragheb, M., Tabakh, H., Pollock, A.J., Woodward, J.J., Dreifus, J.E. and Merrikh, H. (2017) Replication-Transcription Conflicts Generate R-Loops that Orchestrate Bacterial Stress Survival and Pathogenesis. Cell, 170, 787–799 e718.

30. Hamperl, S., Bocek, M.J., Saldivar, J.C., Swigut, T. and Cimprich, K.A. (2017) Transcription-Replication Conflict Orientation Modulates R-Loop Levels and Activates Distinct DNA Damage Responses. Cell, 170, 774–786 e719.

31. Merrikh, H. (2017) Spatial and Temporal Control of Evolution through Replication-Transcription Conflicts. Trends Microbiol, 25, 515–521.

32. Zeman, M.K. and Cimprich, K.A. (2014) Causes and consequences of replication stress. Nat Cell Biol, 16, 2–9.

33. Hull, R.M., King, M., Pizza, G., Krueger, F., Vergara, X. and Houseley, J. (2019) Transcription-induced formation of extrachromosomal DNA during yeast ageing. PLoS Biol, 17, e3000471.

34. Hull, R.M., Cruz, C., Jack, C.V. and Houseley, J. (2017) Environmental change drives accelerated adaptation through stimulated copy number variation. PLoS Biol, 15, e2001333.

35. Zhao, Y., Dominska, M., Petrova, A., Bagshaw, H., Kokoska, R.J. and Petes, T.D. (2017) Properties of Mitotic and Meiotic Recombination in the Tandemly-Repeated CUP1 Gene Cluster in the Yeast Saccharomyces cerevisiae. Genetics, 206, 785–800.

36. Adamo, G.M., Lotti, M., Tamas, M.J. and Brocca, S. (2012) Amplification of the CUP1 gene is associated with evolution of copper tolerance in Saccharomyces cerevisiae. Microbiology (Reading), 158, 2325–2335.

37. Fogel, S., Welch, J.W., Cathala, G. and Karin, M. (1983) Gene amplification in yeast: CUP1 copy number regulates copper resistance. Curr Genet, 7, 347–355.

38. Fogel, S. and Welch, J.W. (1982) Tandem gene amplification mediates copper resistance in yeast. Proc Natl Acad Sci U S A, 79, 5342–5346.

39. Garcia-Muse, T. and Aguilera, A. (2016) Transcription-replication conflicts: how they occur and how they are resolved. Nat Rev Mol Cell Biol, 17, 553–563.

40. Helmrich, A., Ballarino, M., Nudler, E. and Tora, L. (2013) Transcription-replication encounters, consequences and genomic instability. Nat Struct Mol Biol, 20, 412–418.

41. Bermejo, R., Capra, T., Jossen, R., Colosio, A., Frattini, C., Carotenuto, W., Cocito, A., Doksani, Y., Klein, H., Gomez-Gonzalez, B. et al. (2011) The replication checkpoint protects fork stability by releasing transcribed genes from nuclear pores. Cell, 146, 233–246.

42. Bermejo, R., Lai, M.S. and Foiani, M. (2012) Preventing replication stress to maintain genome stability: resolving conflicts between replication and transcription. Mol Cell, 45, 710–718.

43. Li, X. and Manley, J.L. (2005) Inactivation of the SR protein splicing factor ASF/SF2 results in genomic instability. Cell, 122, 365–378.

44. Wang, G. and Vasquez, K.M. (2017) Effects of Replication and Transcription on DNA Structure-Related Genetic Instability. Genes (Basel), 8.

45. Kim, N. and Jinks-Robertson, S. (2011) Guanine repeat-containing sequences confer transcription-dependent instability in an orientation-specific manner in yeast. DNA Repair (Amst), 10, 953–960.

46. Cruz, C. and Houseley, J. (2014) Endogenous RNA interference is driven by copy number. Elife, 3, e01581.

47. Szklarczyk, D., Gable, A.L., Lyon, D., Junge, A., Wyder, S., Huerta-Cepas, J., Simonovic, M., Doncheva, N.T., Morris, J.H., Bork, P. et al. (2019) STRING v11: protein-protein association networks with increased coverage, supporting functional discovery in genome-wide experimental datasets. Nucleic Acids Res, 47, D607–D613.

48. Nepusz, T., Yu, H. and Paccanaro, A. (2012) Detecting overlapping protein complexes in protein-protein interaction networks. Nat Methods, 9, 471–472.

49. Kara, N., Krueger, F., Rugg-Gunn, P. and Houseley, J. (2020) Genome-wide analysis of DNA replication and DNA double strand breaks by TrAEL-seq. bioRxiv

50. Robinson, M.D., McCarthy, D.J. and Smyth, G.K. (2010) edgeR: a Bioconductor package for differential expression analysis of digital gene expression data. Bioinformatics, 26, 139–140.

51. Giaever, G., Chu, A.M., Ni, L., Connelly, C., Riles, L., Veronneau, S., Dow, S., Lucau-Danila, A., Anderson, K., Andre, B. et al. (2002) Functional profiling of the Saccharomyces cerevisiae genome. Nature, 418, 387–391.

52. Smith, C.E., Lam, A.F. and Symington, L.S. (2009) Aberrant double-strand break repair resulting in half crossovers in mutants defective for Rad51 or the DNA polymerase delta complex. Mol Cell Biol, 29, 1432–1441.

53. Che, J., Smith, S., Kim, Y.J., Shim, E.Y., Myung, K. and Lee, S.E. (2015) Hyper-Acetylation of Histone H3K56 Limits Break-Induced Replication by Inhibiting Extensive Repair Synthesis. PLoS Genet, 11, e1004990.

54. Chappidi, N., Nascakova, Z., Boleslavska, B., Zellweger, R., Isik, E., Andrs, M., Menon, S., Dobrovolna, J., Balbo Pogliano, C., Matos, J. et al. (2020) Fork Cleavage-Religation Cycle and Active Transcription Mediate Replication Restart after Fork Stalling at Co-transcriptional R-Loops. Mol Cell, 77, 528–541 e528.

55. Maffia, A., Ranise, C. and Sabbioneda, S. (2020) From R-Loops to G-Quadruplexes: Emerging New Threats for the Replication Fork. Int J Mol Sci, 21.

56. Huertas, P. and Aguilera, A. (2003) Cotranscriptionally formed DNA:RNA hybrids mediate transcription elongation impairment and transcription-associated recombination. Mol Cell, 12, 711–721.

57. Pfeiffer, V., Crittin, J., Grolimund, L. and Lingner, J. (2013) The THO complex component Thp2 counteracts telomeric R-loops and telomere shortening. EMBO J, 32, 2861–2871.

58. Wellinger, R.E., Prado, F. and Aguilera, A. (2006) Replication fork progression is impaired by transcription in hyperrecombinant yeast cells lacking a functional THO complex. Mol Cell Biol, 26, 3327–3334.

59. Hartono, S.R., Malapert, A., Legros, P., Bernard, P., Chedin, F. and Vanoosthuyse, V. (2018) The Affinity of the S9.6 Antibody for Double-Stranded RNAs Impacts the Accurate Mapping of R-Loops in Fission Yeast. J Mol Biol, 430, 272–284.

60. Lockhart, A., Pires, V.B., Bento, F., Kellner, V., Luke-Glaser, S., Yakoub, G., Ulrich, H.D. and Luke, B. (2019) RNase H1 and H2 Are Differentially Regulated to Process RNA-DNA Hybrids. Cell Rep, 29, 2890–2900 e2895.

61. Chang, E.Y., Tsai, S., Aristizabal, M.J., Wells, J.P., Coulombe, Y., Busatto, F.F., Chan, Y.A., Kumar, A., Dan Zhu, Y., Wang, A.Y. et al. (2019) MRE11-RAD50-NBS1 promotes Fanconi Anemia R-loop suppression at transcription-replication conflicts. Nat Commun, 10, 4265.

62. Ellisdon, A.M., Dimitrova, L., Hurt, E. and Stewart, M. (2012) Structural basis for the assembly and nucleic acid binding of the TREX-2 transcription-export complex. Nat Struct Mol Biol, 19, 328–336.

63. Gonzalez-Aguilera, C., Tous, C., Gomez-Gonzalez, B., Huertas, P., Luna, R. and Aguilera, A. (2008) The THP1-SAC3-SUS1-CDC31 complex works in transcription elongation-mRNA export preventing RNA-mediated genome instability. Mol Biol Cell, 19, 4310–4318.

64. Schneider, M., Hellerschmied, D., Schubert, T., Amlacher, S., Vinayachandran, V., Reja, R., Pugh, B.F., Clausen, T. and Kohler, A. (2015) The Nuclear Pore-Associated TREX-2 Complex Employs Mediator to Regulate Gene Expression. Cell, 162, 1016–1028.

65. Santos-Pereira, J.M., Garcia-Rubio, M.L., Gonzalez-Aguilera, C., Luna, R. and Aguilera, A. (2014) A genome-wide function of THSC/TREX-2 at active genes prevents transcription-replication collisions. Nucleic Acids Res, 42, 12000–12014.

66. Verger, A., Monte, D. and Villeret, V. (2019) Twenty years of Mediator complex structural studies. Biochem Soc Trans, 47, 399–410.

67. Lariviere, L., Plaschka, C., Seizl, M., Petrotchenko, E.V., Wenzeck, L., Borchers, C.H. and Cramer, P. (2013) Model of the Mediator middle module based on protein cross-linking. Nucleic Acids Res, 41, 9266–9273.

68. Guglielmi, B., van Berkum, N.L., Klapholz, B., Bijma, T., Boube, M., Boschiero, C., Bourbon, H.M., Holstege, F.C. and Werner, M. (2004) A high resolution protein interaction map of the yeast Mediator complex. Nucleic Acids Res, 32, 5379–5391.

69. Lariviere, L., Geiger, S., Hoeppner, S., Rother, S., Strasser, K. and Cramer, P. (2006) Structure and TBP binding of the Mediator head subcomplex Med8-Med18-Med20. Nat Struct Mol Biol, 13, 895–901.

70. Schilbach, S., Hantsche, M., Tegunov, D., Dienemann, C., Wigge, C., Urlaub, H. and Cramer, P. (2017) Structures of transcription pre-initiation complex with TFIIH and Mediator. Nature, 551, 204–209.

71. Casamassimi, A. and Napoli, C. (2007) Mediator complexes and eukaryotic transcription regulation: an overview. Biochimie, 89, 1439–1446.

72. Lei, E.P., Stern, C.A., Fahrenkrog, B., Krebber, H., Moy, T.I., Aebi, U. and Silver, P.A. (2003) Sac3 is an mRNA export factor that localizes to cytoplasmic fibrils of nuclear pore complex. Mol Biol Cell, 14, 836–847.

73. Dieppois, G. and Stutz, F. (2010) Connecting the transcription site to the nuclear pore: a multi-tether process that regulates gene expression. J Cell Sci, 123, 1989–1999.

74. Ahmed, S. and Brickner, J.H. (2007) Regulation and epigenetic control of transcription at the nuclear periphery. Trends Genet, 23, 396–402.

75. Blobel, G. (1985) Gene gating: a hypothesis. Proc Natl Acad Sci U S A, 82, 8527–8529.

76. Liang, Q. and Zhou, B. (2007) Copper and manganese induce yeast apoptosis via different pathways. Mol Biol Cell, 18, 4741–4749.

77. Rass, U. (2013) Resolving branched DNA intermediates with structure-specific nucleases during replication in eukaryotes. Chromosoma, 122, 499–515.

78. Wyatt, H.D. and West, S.C. (2014) Holliday junction resolvases. Cold Spring Harb Perspect Biol, 6, a023192.

79. Pardo, B., Moriel-Carretero, M., Vicat, T., Aguilera, A. and Pasero, P. (2020) Homologous recombination and Mus81 promote replication completion in response to replication fork blockage. EMBO Rep, 21, e49367.

80. Osborn, A.J. and Elledge, S.J. (2003) Mrc1 is a replication fork component whose phosphorylation in response to DNA replication stress activates Rad53. Genes Dev, 17, 1755–1767.

81. Alcasabas, A.A., Osborn, A.J., Bachant, J., Hu, F., Werler, P.J., Bousset, K., Furuya, K., Diffley, J.F., Carr, A.M. and Elledge, S.J. (2001) Mrc1 transduces signals of DNA replication stress to activate Rad53. Nat Cell Biol, 3, 958–965.

82. Pardo, B., Crabbe, L. and Pasero, P. (2017) Signaling pathways of replication stress in yeast. FEMS Yeast Res, 17.

83. Zhao, X., Georgieva, B., Chabes, A., Domkin, V., Ippel, J.H., Schleucher, J., Wijmenga, S., Thelander, L. and Rothstein, R. (2000) Mutational and structural analyses of the ribonucleotide reductase inhibitor Sml1 define its Rnr1 interaction domain whose inactivation allows suppression of mec1 and rad53 lethality. Mol Cell Biol, 20, 9076–9083.

84. Shimada, K., Pasero, P. and Gasser, S.M. (2002) ORC and the intra-S-phase checkpoint: a threshold regulates Rad53p activation in S phase. Genes Dev, 16, 3236–3252.

85. Somyajit, K., Gupta, R., Sedlackova, H., Neelsen, K.J., Ochs, F., Rask, M.B., Choudhary, C. and Lukas, J. (2017) Redox-sensitive alteration of replisome architecture safeguards genome integrity. Science, 358, 797–802.

86. Yeeles, J.T.P., Janska, A., Early, A. and Diffley, J.F.X. (2017) How the Eukaryotic Replisome Achieves Rapid and Efficient DNA Replication. Mol Cell, 65, 105–116.

87. Calzada, A., Hodgson, B., Kanemaki, M., Bueno, A. and Labib, K. (2005) Molecular anatomy and regulation of a stable replisome at a paused eukaryotic DNA replication fork. Genes Dev, 19, 1905–1919.

88. Tourriere, H., Versini, G., Cordon-Preciado, V., Alabert, C. and Pasero, P. (2005) Mrc1 and Tof1 promote replication fork progression and recovery independently of Rad53. Mol Cell, 19, 699–706.

89. Minocherhomji, S., Ying, S., Bjerregaard, V.A., Bursomanno, S., Aleliunaite, A., Wu, W., Mankouri, H.W., Shen, H., Liu, Y. and Hickson, I.D. (2015) Replication stress activates DNA repair synthesis in mitosis. Nature, 528, 286–290.

90. McCune, H.J., Danielson, L.S., Alvino, G.M., Collingwood, D., Delrow, J.J., Fangman, W.L., Brewer, B.J. and Raghuraman, M.K. (2008) The temporal program of chromosome replication: genomewide replication in clb5{Delta} Saccharomyces cerevisiae. Genetics, 180, 1833–1847.

91. Schwob, E. and Nasmyth, K. (1993) CLB5 and CLB6, a new pair of B cyclins involved in DNA replication in Saccharomyces cerevisiae. Genes Dev, 7, 1160–1175.

92. Donaldson, A.D., Raghuraman, M.K., Friedman, K.L., Cross, F.R., Brewer, B.J. and Fangman, W.L. (1998) CLB5-dependent activation of late replication origins in S. cerevisiae. Mol Cell, 2, 173–182.

93. Koren, A., Soifer, I. and Barkai, N. (2010) MRC1-dependent scaling of the budding yeast DNA replication timing program. Genome Res, 20, 781–790.

94. Dovrat, D., Dahan, D., Sherman, S., Tsirkas, I., Elia, N. and Aharoni, A. (2018) A Live-Cell Imaging Approach for Measuring DNA Replication Rates. Cell Rep, 24, 252–258.

95. Loeillet, S., Herzog, M., Puddu, F., Legoix, P., Baulande, S., Jackson, S.P. and Nicolas, A.G. (2020) Trajectory and uniqueness of mutational signatures in yeast mutators. Proc Natl Acad Sci U S A, 117, 24947–24956.

96. Omer, S., Lavi, B., Mieczkowski, P.A., Covo, S. and Hazkani-Covo, E. (2017) Whole Genome Sequence Analysis of Mutations Accumulated in rad27Delta Yeast Strains with Defects in the Processing of Okazaki Fragments Indicates Template-Switching Events. G3 (Bethesda), 7, 3775–3787.

97. Lengronne, A. and Schwob, E. (2002) The yeast CDK inhibitor Sic1 prevents genomic instability by promoting replication origin licensing in late G(1). Mol Cell, 9, 1067–1078.

98. Liachko, I., Bhaskar, A., Lee, C., Chung, S.C., Tye, B.K. and Keich, U. (2010) A comprehensive genome-wide map of autonomously replicating sequences in a naive genome. PLoS Genet, 6, e1000946.

99. Wyrick, J.J., Aparicio, J.G., Chen, T., Barnett, J.D., Jennings, E.G., Young, R.A., Bell, S.P. and Aparicio, O.M. (2001) Genome-wide distribution of ORC and MCM proteins in S. cerevisiae: high-resolution mapping of replication origins. Science, 294, 2357–2360.

100. Kramarz, K., Schirmeisen, K., Boucherit, V., Ait Saada, A., Lovo, C., Palancade, B., Freudenreich, C. and Lambert, S.A.E. (2020) The nuclear pore primes recombination-dependent DNA synthesis at arrested forks by promoting SUMO removal. Nat Commun, 11, 5643.

101. Garcia-Benitez, F., Gaillard, H. and Aguilera, A. (2017) Physical proximity of chromatin to nuclear pores prevents harmful R loop accumulation contributing to maintain genome stability. Proc Natl Acad Sci U S A, 114, 10942–10947.

102. Doi, G., Okada, S., Yasukawa, T., Sugiyama, Y., Bala, S., Miyazaki, S., Kang, D. and Ito, T. (2021) Catalytically inactive Cas9 impairs DNA replication fork progression to induce focal genomic instability. Nucleic Acids Res, 49, 954–968.

103. Cipponi, A., Goode, D.L., Bedo, J., McCabe, M.J., Pajic, M., Croucher, D.R., Rajal, A.G., Junankar, S.R., Saunders, D.N., Lobachevsky, P. et al. (2020) MTOR signaling orchestrates stress-induced mutagenesis, facilitating adaptive evolution in cancer. Science, 368, 1127–1131.

104. Russo, M., Crisafulli, G., Sogari, A., Reilly, N.M., Arena, S., Lamba, S., Bartolini, A., Amodio, V., Magri, A., Novara, L. et al. (2019) Adaptive mutability of colorectal cancers in response to targeted therapies. Science, 366, 1473–1480.

105. Foster, P.L. (2007) Stress-induced mutagenesis in bacteria. Crit Rev Biochem Mol Biol, 42, 373–397.

106. Fitzgerald, D.M., Hastings, P.J. and Rosenberg, S.M. (2017) Stress-Induced Mutagenesis: Implications in Cancer and Drug Resistance. Annu Rev Cancer Biol, 1, 119–140.

107. Merrikh, C.N. and Merrikh, H. (2018) Gene inversion potentiates bacterial evolvability and virulence. Nat Commun, 9, 4662.

108. Macheret, M., Bhowmick, R., Sobkowiak, K., Padayachy, L., Mailler, J., Hickson, I.D. and Halazonetis, T.D. (2020) High-resolution mapping of mitotic DNA synthesis regions and common fragile sites in the human genome through direct sequencing. Cell Res, 30, 997–1008.

109. Sarni, D., Sasaki, T., Miron, K., Tur-Sinai, M.I., Rivera-Mulia, J.C., Magnuson, B., Lungman, M., Gilbert, D.M. and Kerem, B. (2019) Replication Timing and Transcription Identifies a Novel Fragility Signature Under Replication Stress. BioRxiv.

110. Brison, O., El-Hilali, S., Azar, D., Koundrioukoff, S., Schmidt, M., Nahse, V., Jaszczyszyn, Y., Lachages, A.M., Dutrillaux, B., Thermes, C. et al. (2019) Transcription-mediated organization of the replication initiation program across large genes sets common fragile sites genome-wide. Nat Commun, 10, 5693.

111. Debatisse, M., Le Tallec, B., Letessier, A., Dutrillaux, B. and Brison, O. (2012) Common fragile sites: mechanisms of instability revisited. Trends Genet, 28, 22–32.

112. Durkin, S.G. and Glover, T.W. (2007) Chromosome fragile sites. Annu Rev Genet, 41, 169–192.

113. Le Beau, M.M., Rassool, F.V., Neilly, M.E., Espinosa, R., 3rd, Glover, T.W., Smith, D.I. and McKeithan, T.W. (1998) Replication of a common fragile site, FRA3B, occurs late in S phase and is delayed further upon induction: implications for the mechanism of fragile site induction. Hum Mol Genet, 7, 755–761.

114. Fisher, M.C., Hawkins, N.J., Sanglard, D. and Gurr, S.J. (2018) Worldwide emergence of resistance to antifungal drugs challenges human health and food security. Science, 360, 739–742.

115. Fisher, M.C., Garner, T.W. and Walker, S.F. (2009) Global emergence of Batrachochytrium dendrobatidis and amphibian chytridiomycosis in space, time, and host. Annu Rev Microbiol, 63, 291–310.

116. Xu, W., Aparicio, J.G., Aparicio, O.M. and Tavare, S. (2006) Genome-wide mapping of ORC and Mcm2p binding sites on tiling arrays and identification of essential ARS consensus sequences in S. cerevisiae. BMC Genomics, 7, 276.

