## Supplementary figures for "Stimulation of adaptive gene amplification by origin firing under replication fork constraint"

### Supplementary Material

#### Supplementary Figures

Supplementary figures are numbered 2-6 as each forms a supplement to one of the main figures 2-6.

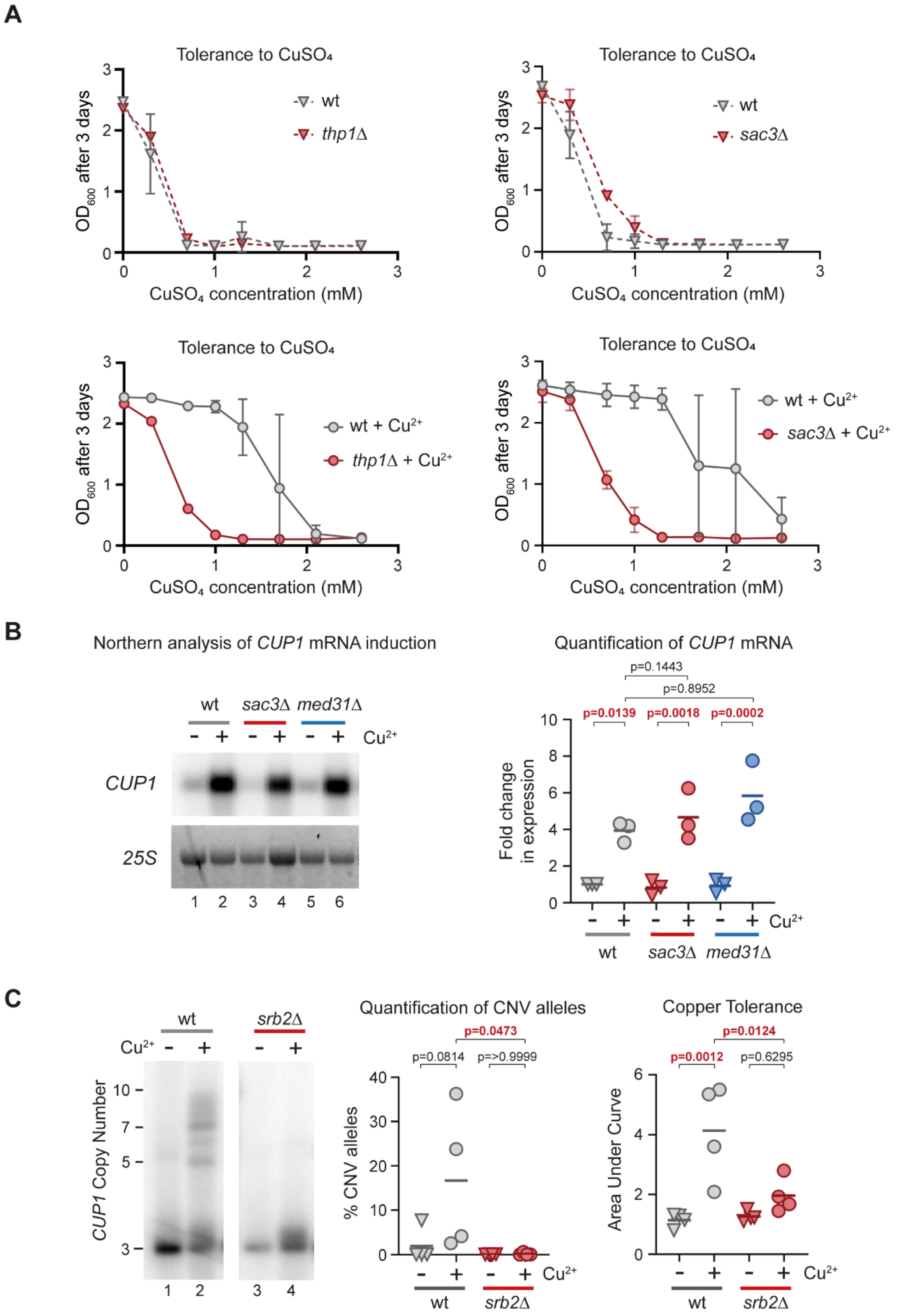

**Supplementary Figure 2.** Supplement to TREX-2 and Mediator are required for transcription stimulated *CUP1* CNV. **(A)** Plots of final $OD_{660}$ versus [$CuSO_4$] for wild type, *thp1*Δ and *sac3*Δ. Upper plots show naïve cells that have not been pre-exposed to $CuSO_4$, lower plots show cells that have been pre-cultured in 0.3mM $CuSO_4$. Tolerance of mutants that have not been pre-cultured in $CuSO_4$ is similar or higher than wild type, showing that loss of Thp1 and Sac3 does not impair the normal response to environmental copper. This is the source data underlying the plots in Figures 2B, 2C. **(B)** Northern blot analysis of *CUP1* mRNA, in wild-type, *sac3*Δ and *med31*Δ cells. Mid-log cells were treated with 0.3mM $CuSO_4$ for 6 hours and fold change in *CUP1* expression was quantified relative to the *25S* rRNA highlighted by ethidium bromide staining, p-values calculated by 1-way ANOVA; n = 3. **(C)** Southern blot analysis of *CUP1* copy number and copper tolerance analysis of *3xCUP1* wild-type (wt) and *srb2*Δ cells, performed as in Figure 2A, n = 3.

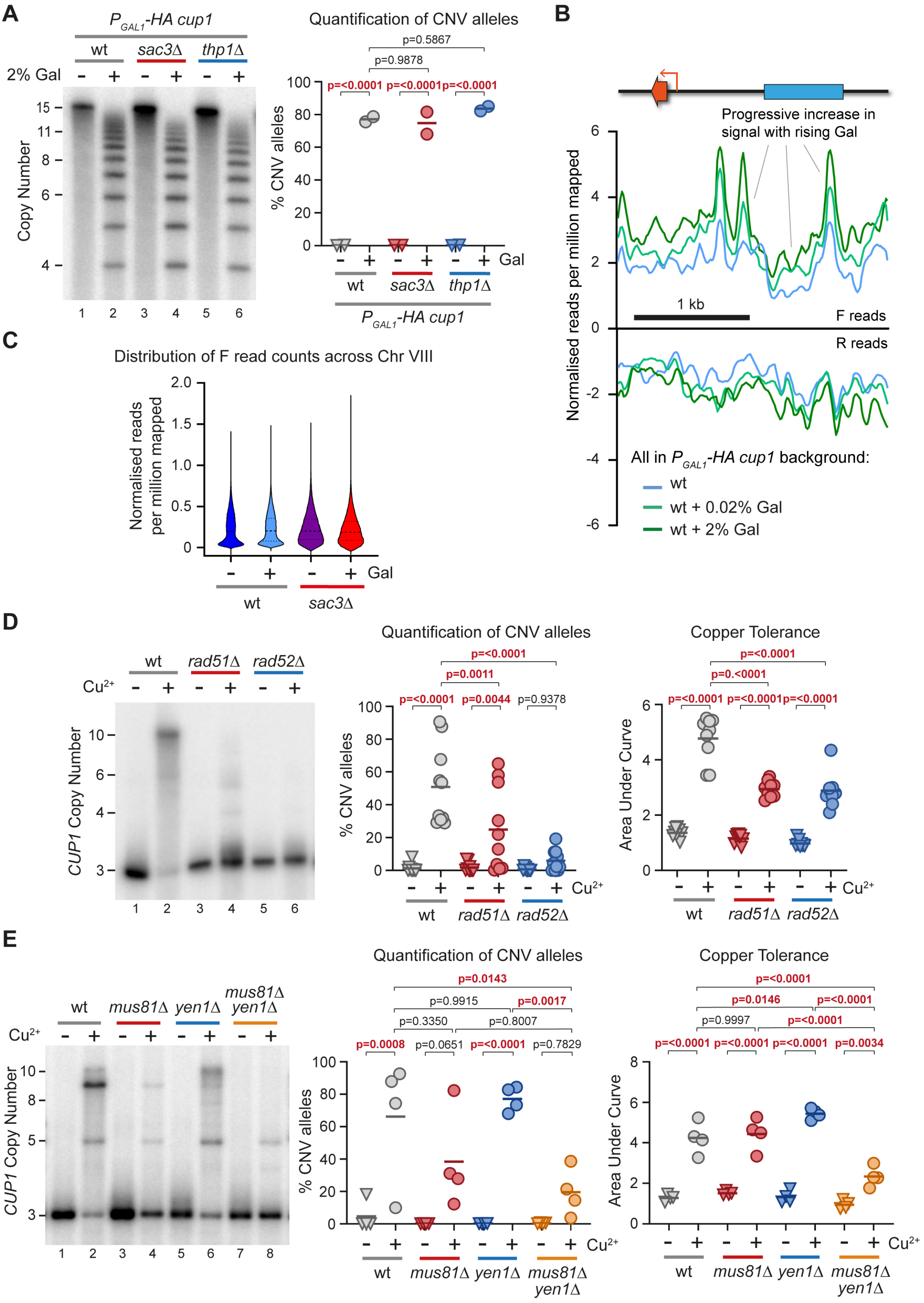
A
$P_{GAL1}$-HA cup1
wt sac3Δ thp1Δ
2% Gal
Copy Number
Quantification of CNV alleles
p=0.5867
p=0.9878
p=<0.0001 p=<0.0001 p=<0.0001
% CNV alleles
Gal
B
Progressive increase in signal with rising Gal
Normalised reads per million mapped
1 kb
F reads
R reads
All in $P_{GAL1}$-HA cup1 background:
wt
wt + 0.02% Gal
wt + 2% Gal
C
Distribution of F read counts across Chr VIII
Normalised reads per million mapped
D
wt rad51Δ rad52Δ
$Cu^{2+}$
CUP1 Copy Number
Quantification of CNV alleles
p=<0.0001
p=0.0011
p=<0.0001 p=0.0044 p=0.9378
Copper Tolerance
p=<0.0001
p=0.<0001
p=<0.0001 p=<0.0001 p=<0.0001
Area Under Curve
E
wt mus81Δ yen1Δ mus81Δ yen1Δ
p=0.0143
p=0.9915 p=0.0017
p=0.3350 p=0.8007
p=0.0008 p=0.0651 p=<0.0001 p=0.7829
p=<0.0001
p=0.0146 p=<0.0001
p=0.9997 p=<0.0001
p=<0.0001 p=<0.0001 p=<0.0001 p=0.0034

**Supplementary Figure 3.** Supplement to replication fork stalling and cleavage at *CUP1* locus. **(A)** Cells with every *CUP1* ORF and promoter in each *CUP1* copy replaced by $P_{GAL1}$-*HA*, grown for 10 generations in raffinose ± 2% galactose. Analysis by Southern blot comparing *sac3*Δ and *thp1*Δ cells to wild type. Quantification shows the percentage of alleles deviating from the parental copy number of 17 copies; n = 2. **(B)** Plots of TrAEL-seq read density on forward and reverse strands across a $P_{GAL1}$-*HA cup1* repeat for wild-type cells induced for 6 hours with 0%, 0.02% or 2% galactose. TrAEL-seq profiles are an average of two biological replicates, 0% and 0.02% conditions are the same data as Figure 3C, data is processed as in Figure 3C. **(C)** Violin plots showing the distribution of read counts for all 50 bp windows spaced at 10 bp intervals across chromosome VIII that do not overlap multi-copy elements, data was processed as in Figure 3C. **(D)** and **(E)** Southern blot analysis of *CUP1* copy number and copper adaptation analysis of *3xCUP1* wild-type (wt) and indicated mutant cells, performed as Figure 3E, n = 10 for (D), n = 4 for (E)**.**

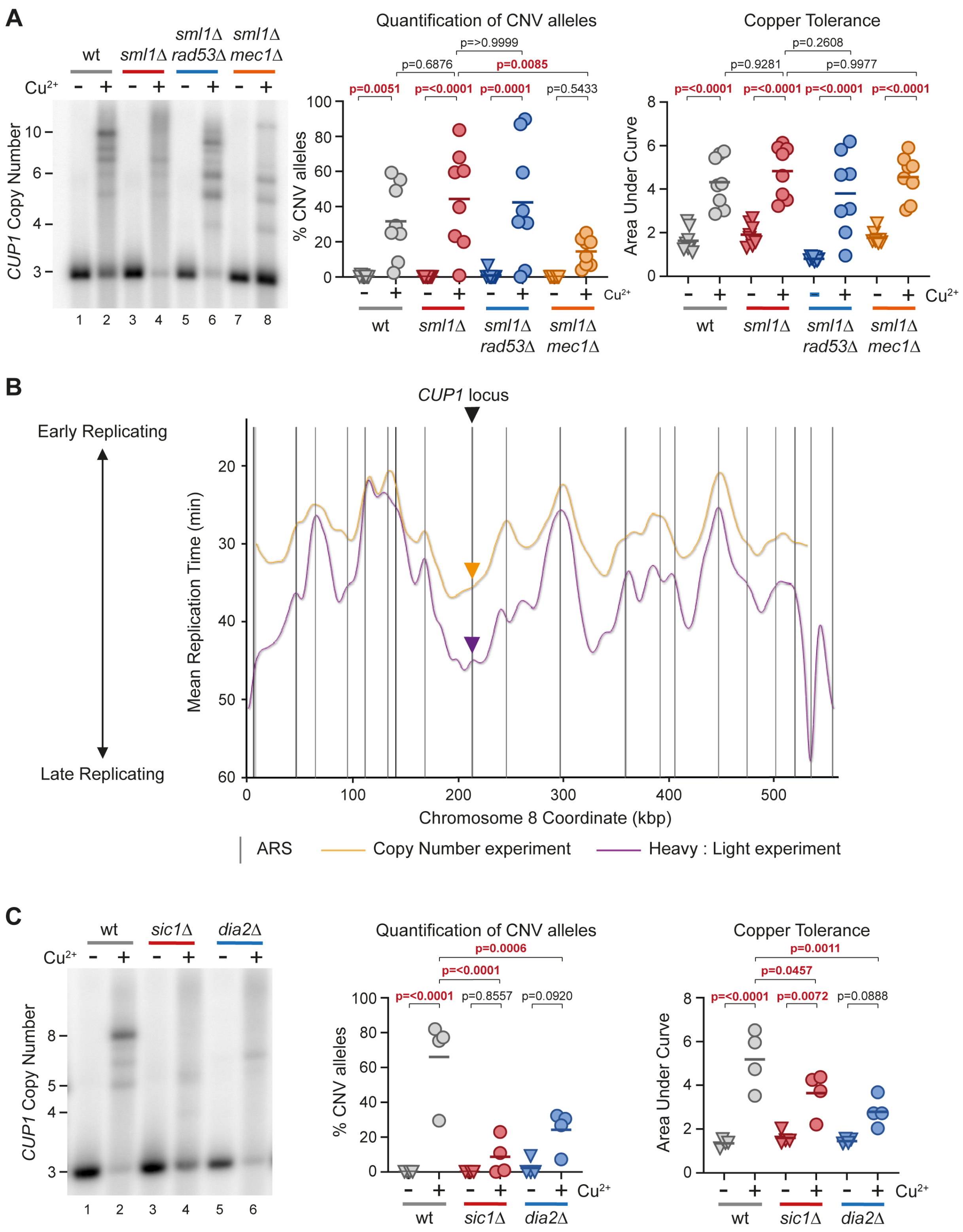


**Supplementary Figure 4.** Supplement to replication timing and fork progression control *CUP1* CNV. **(A)** Southern blot analysis of *CUP1* copy number and copper adaptation analysis of *3xCUP1* wild-type (wt), *sml1Δ*, *rad53Δ sml1Δ* and *mec1Δ sml1Δ* cells, performed as in Figure 4A, n = 8. **(B)** Replication timing on Chromosome 8 from OriDB (1), with data collected by two independent studies measuring change in copy number (2) (shown in yellow) or Heavy : Light isotope transfer (3) (shown in purple). Grey lines correspond to Replication origins and arrows indicate the late replication timing of the *CUP1* locus from the respective studies. **(C)** Southern and growth curve analysis of *3xCUP1* wildtype (wt), and indicated mutant grown for 10 generations ± 0.3 mM $CuSO_4$ as in Figure 4A; n = 4.

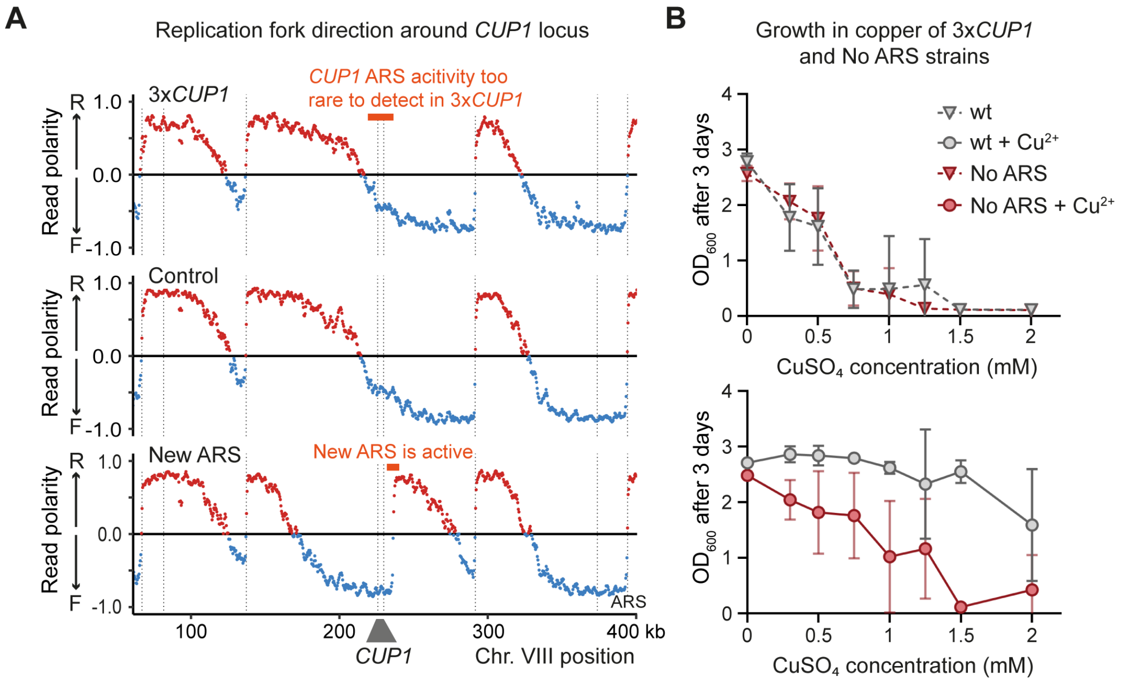


**Supplementary Figure 5.** Supplement to local replication origin firing regulates CNV at *CUP1* locus. **(A)** TrAEL-seq read polarity plots for 3x*CUP1* strains wild type, control or New ARS (defined in Figure 5B), showing region surrounding *CUP1* on chromosome VIII, processed as Figure 5A, n=1. **(B)** Plots of final $OD_{660}$ versus [$CuSO_4$] for wild type and No ARS cells that are naïve (above) or pre-cultured in 0.3mM $CuSO_4$ (below). Copper resistance of No ARS cells that have not been pre-cultured in $CuSO_4$ is similar to wild type, showing that deletion of this large region of the *CUP1* repeat does not impair the normal response to environmental copper. This is the source data underlying the plots in Figure 5D.

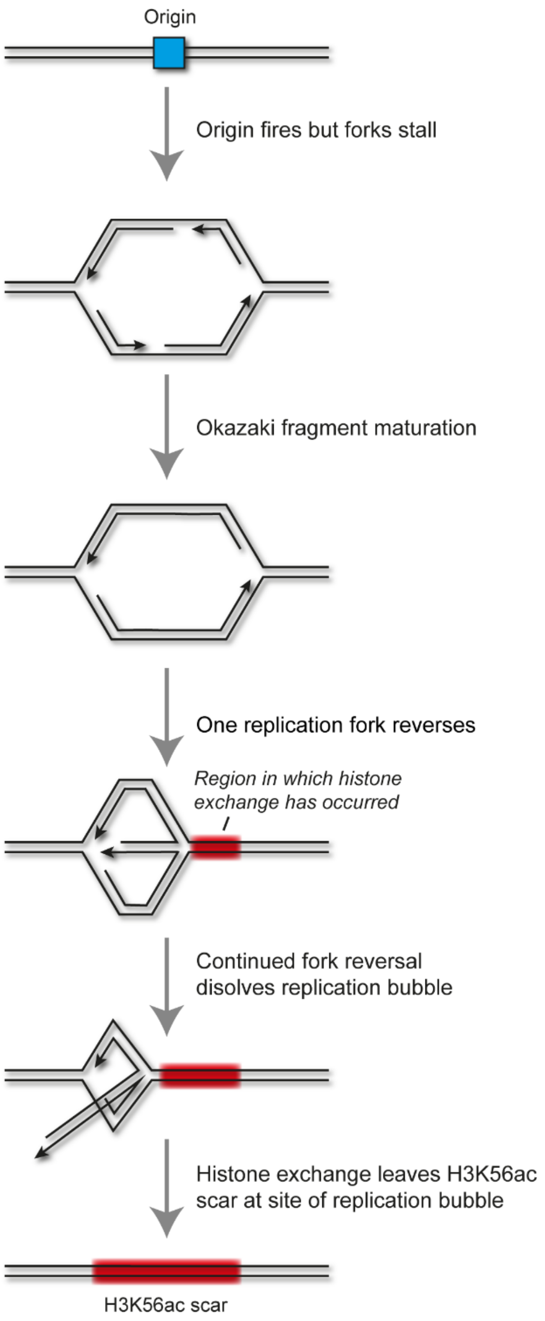


**Supplementary Figure 6.** Proposed mechanism for H3K56ac scar incorporation during unsuccessful replication origin firing. DNA strands are shown in black, the replication origin is shown in blue and chromatin containing H3K56ac shown in red. In the presence of topological stress, replication origins may fire but stall soon after. Positive supercoiling may promote reversal of such stalled fork structures to the point of dissolution, leaving behind a “H3K56ac scar” from initial chromatin assembly

**Supplementary Tables**

Tables are provided as Excel files.

**S1 Table** *S. cerevisiae* yeast strains used in this study

**S2 Table** Oligonucleotide pairs used in this study.

**S3 Table** Quantification of CNV derived from Mutant Screen. % CNV alleles were calculated as intensity of bands by southern blot analysis: CNV alleles / (CNV alleles + Parental allele) x 100. Fold-change in CNV was calculated as % CNV alleles for *gene*Δ / % CNV alleles of Wild Type.

**Supplementary References**

1. Siow, C.C., Nieduszynska, S.R., Muller, C.A. and Nieduszynski, C.A. (2012) OriDB, the DNA replication origin database updated and extended. *Nucleic Acids Res*, **40**, D682-686.
2. Yabuki, N., Terashima, H. and Kitada, K. (2002) Mapping of early firing origins on a replication profile of budding yeast. *Genes Cells*, **7**, 781-789.
3. Raghuraman, M.K., Winzeler, E.A., Collingwood, D., Hunt, S., Wodicka, L., Conway, A., Lockhart, D.J., Davis, R.W., Brewer, B.J. and Fangman, W.L. (2001) Replication dynamics of the yeast genome. *Science*, **294**, 115-121.
4. Brachmann, C.B., Davies, A., Cost, G.J., Caputo, E., Li, J., Hieter, P. and Boeke, J.D. (1998) Designer deletion strains derived from Saccharomyces cerevisiae S288C: a useful set of strains and plasmids for PCR-mediated gene disruption and other applications. *Yeast*, **14**, 115-132.
5. Dai, J., Hyland, E.M., Yuan, D.S., Huang, H., Bader, J.S. and Boeke, J.D. (2008) Probing nucleosome function: a highly versatile library of synthetic histone H3 and H4 mutants. *Cell*, **134**, 1066-1078.
6. Breslow, D.K., Cameron, D.M., Collins, S.R., Schuldiner, M., Stewart-Ornstein, J., Newman, H.W., Braun, S., Madhani, H.D., Krogan, N.J. and Weissman, J.S. (2008) A comprehensive strategy enabling high-resolution functional analysis of the yeast genome. *Nat Methods*, **5**, 711-718.
7. Hull, R.M., Cruz, C., Jack, C.V. and Houseley, J. (2017) Environmental change drives accelerated adaptation through stimulated copy number variation. *PLoS Biol*, **15**, e2001333.
